# Hypothalamic-pituitary-adrenal axis dysfunction worsens epilepsy outcomes and increases SUDEP risk

**DOI:** 10.1101/2022.03.15.484525

**Authors:** Trina Basu, Pantelis Antonoudiou, Grant L. Weiss, Daniel Friedman, Juliana Laze, Orrin Devinsky, Jamie Maguire

## Abstract

Epilepsy is often comorbid with psychiatric illnesses, including anxiety and depression. Despite the high incidence of psychiatric comorbidities in people with epilepsy, few studies address the underlying mechanisms. Stress can trigger both epilepsy and depression and significant accumulating evidence, from both human and animal studies, support that hypothalamic-pituitary-adrenal (HPA) axis dysfunction may contribute to the comorbidity of these two disorders^1^. Here we directly investigate the contribution of HPA axis dysfunction to epilepsy outcomes and psychiatric comorbidities. We generated a novel mouse model (KCC2/Crh mice) that lacks the K^+^/Cl^-^ co-transporter, KCC2, in corticotropin-releasing hormone (CRH) neurons, which exhibit stress- and seizure-induced hyperactivation of the HPA axis.^2,3^ Here we demonstrate that HPA axis dysfunction in KCC2/Crh mice is associated with increased vulnerability to behavioral deficits in chronically epileptic mice and more severe neuropathological features associated with chronic epilepsy (e.g. increased mossy fiber sprouting). Despite equivalent seizure burden, HPA axis dysfunction in these chronically epileptic mice increased the incidence of sudden unexpected death in epilepsy (SUDEP). Suppressing HPA axis hyperexcitability in this model using a pharmacological or chemogenetic approach decreased SUDEP incidence, supporting HPA axis dysfunction in SUDEP vulnerability. We also observed changes in neuroendocrine markers in individuals that died of SUDEP compared to people with epilepsy or individuals without a history of epilepsy. Together, these findings describe a novel, nongenetic mouse model of SUDEP which will benefit preclinical SUDEP research to gain a better understanding of the underlying pathological mechanisms contributing to SUDEP. HPA axis dysfunction increases vulnerability to psychiatric comorbidities in epilepsy and may be a risk factor for SUDEP.

## Introduction

Psychiatric comorbidities are highly prevalent in people with epilepsy (PWE), affecting approximately 75%, with depression (55%) and anxiety (25-50%) being the most common.^4,5^ The comorbid appearance of psychiatric disorders in PWE occurs at rates much higher than both the general population and in people with chronic illness other than epilepsy.^4,6^ Increasing evidence suggests that this comorbidity has neurobiological underpinnings,^7^ but the mechanisms underlying mood disorders in epilepsy remain unclear. Since seizure control remains the primary treatment focus for PWE, psychiatric comorbidities in people with epilepsy are often underdiagnosed and undertreated. As a result, these comorbidities can negatively impact the quality of life for people with epilepsy and, if left untreated, can severely worsen the course of epilepsy and/or increase suicide risk in PWE.^8,9^

A cardinal feature of depression, the psychiatric disorder most diagnosed in PWE, is hypothalamic-pituitary-adrenal (HPA) axis hyperactivity.^10–13^ The HPA axis mediates the body’s physiological response to stress, a major risk factor for depression and anxiety^14^ and a seizure trigger in many PWE.^15,16^ Psychological or physiological stress induces a HPA-mediated a neuroendocrine response. Corticotropin-releasing hormone (CRH) neurons in the paraventricular nucleus (PVN) of the hypothalamus govern HPA axis control. In response to stress or seizures,^2,17^ CRH is released and sequentially triggers the release of adrenocorticotropin hormone (ACTH) from the anterior pituitary gland, and cortisol from the adrenal glands. Cortisol (in humans) and corticosterone (in rodents) exert their effects through actions on either mineralocorticoid receptors or glucocortiocoid receptors. Many people who suffer from depression exhibit HPA axis dysregulation, as evidenced by elevated CRH and cortisol levels.^18,19^ In fact, hypercortisolism is a hallmark feature of major depression.^11,20^

Interestingly, up to 60% of PWE also report stress as a trigger for seizures,^15,21,22^ suggesting that HPA axis dysregulation may drive some elements of seizure susceptibility in epilepsy. In addition, cortisol levels are basally elevated in PWE and are further increased following a seizure, which correlates with seizure severity.^18,19^ Similarly, we found that seizures alone activate the HPA axis in rodents.^2^ Hyperactivation of the HPA axis causes compounding pathology as stress hormones exert proconvulsant actions^23^ and HPA axis dysfunction has been shown to exacerbate neuropathology and disease progression in epilepsy and accelerate epileptogenesis.^17,24–27^ These studies suggest that controlling the body’s physiological response to stress may be an effective treatment for seizure control. Indeed, we have shown that blocking seizure induced elevations of corticosterone with a CRH antagonist, antalarmin, reduces future seizure susceptibility.^2^ We have also demonstrated that blunting HPA axis activation improved outcomes in chronically epileptic mice.^24^ Recent studies also show that attenuating HPA axis activity through use of the glucocorticoid antagonist, RU486, or the glucocorticoid specific inhibitor, CORT108297, mitigates *status epilepticus* (SE) induced neuropathology.^28,29^ However, little is known regarding how seizure induced HPA axis activation contributes to comorbid psychiatric disorders in epilepsy. We recently demonstrated that a mouse model of pilocarpine-induced temporal lobe epilepsy (TLE) exhibits stress-induced elevations in circulating corticosterone and increased anxiety- and depression-like behaviors.^17,24^ Our previous work demonstrated that suppressing the seizure-induced activation of the HPA axis decreased seizure burden and comorbid behavioral deficits.^24^ However, these previous studies rely on correlations between corticosterone levels and epilepsy outcomes. Here, we directly examine the relationship between HPA axis dysfunction and psychiatric comorbidities associated with epilepsy using a genetic mouse model exhibiting HPA axis hyperactivation.

The HPA axis is regulated by GABAergic control of CRH neurons in the PVN.^30,31^ Our lab found that stress- and seizure-induced activation of the HPA axis is driven by a collapse in the chloride gradient in CRH neurons, required for GABAergic inhibition and maintained by the K^+^/Cl^-^ co-transporter, KCC2.^2,32^ To explore the role of HPA axis dysfunction in epilepsy and associated psychiatric comorbidities, we generated mice that lack KCC2 specifically in CRH neurons (*Kcc2/Crh*).^3^ The *Kcc2/Crh* mice exhibit an exaggerated seizure induced activation of the HPA axis. Here, we show that seizure induced hyperactivation of the HPA axis in male *Kcc2/Crh* mice contributes to increased vulnerability to anxiety- and depression-like behaviors and greater neuropathological changes associated with epilepsy. Unexpectedly, HPA dysfunction in chronically epileptic *Kcc2/Crh* mice is also associated with a profound increase in susceptibility to sudden unexpected death in epilepsy (SUDEP). We demonstrate that pharmacological or chemogenetic attenuation of seizure induced activation of the HPA axis reduces seizure burden and SUDEP incidence, further implicating HPA axis dysfunction in SUDEP. While certain genetic risk factors increase SUDEP predisposition,^33,34^ non-genetic influences contributing to SUDEP are not yet understood and there is a lack of experimental models to explore additional pathophysiological mechanisms. Our work suggests that HPA axis dysfunction may increase risk for SUDEP and that the *Kcc2/Crh* mouse model is a novel, non-genetic model of SUDEP with utility for examining non-genetic mechanisms contributing to SUDEP. Blood samples obtained from PWE and PWE with suspected SUDEP indicate that persistent and excessive HPA axis dysfunction may contribute to SUDEP, suggesting that identifying sources of HPA axis dysfunction may serve as novel biomarkers for those at risk of SUDEP.

## Methods

### Animals

We used adult (8-12 weeks) male Cre^-/-^ (WT) and *Kcc2/Crh* mice bred on a 129/Sv background strain and housed at the Tufts University School of Medicine’s Division of Laboratory Animal Medicine facility. Food and water were provided *ad libitum*. All procedures used in this study were approved by the Tufts University Institutional Animal Care and Use Committee. Our lab has generated and has previously characterized the *Kcc2/Crh* mouse line.^3^ Briefly, we crossed floxed KCC2 (*Kcc2^f/f^*) mice (a generous gift from Dr. Stephen Moss) with CRH-Cre mice obtained from the Mutant Mouse Regional Research Center (Stock #030850-UCD) to generate mice that lack KCC2 in CRH neurons (*Kcc2/Crh*).^3^ We genotyped the *Kcc2/Crh* mice in house and through Transnetyx using the following primers:

KCC2
5′: ATGAGTAGCAGATCCCATAGGCGAACC
3′ CTGCCAAGAGCCATTACTACAGTGGATG

CRH-Cre
5′: CTGTCTTGTCTGTGGGTGTCCGAT
3′: CGGCAAACGGACAGAAGCATT

The expected polymerase chain reaction (PCR) product size for the floxed KCC2 mice is 543 bp and 426 bp for the WT mice.

### Ventral intrahippocampal kainic acid injections

We followed the ventral intrahippocampal kainic acid (vIHKA) protocol previously reported by Zeidler et al., 2018.^35^ Mice were first anesthetized with 100 mg/kg dose of ketamine and 10 mg/kg dose of xylazine and administered 0.5 mg/mL dose of Buprenorphine Sustained Release Lab formulation preoperatively. Kainic acid (100 nL of 20 mM; vIHKA) or vehicle (100 nL of sterile saline; vIHSA) was stereotaxically injected into the ventral hippocampus: 3.6 mm posterior, −2.8 mm lateral, and 2.8 mm depth. Seizures were not pharmacologically terminated and only mice that achieved *status epilepticus* were used for this study.

### Gi DREADD injections and CNO administration

A subset of mice were bilaterally injected with AAV-hSyn-DIO-hM4D(Gi)-mCherry (Gi DREADD) in the PVN of the hypothalamus in *Kcc2/Crh* mice, an approach which has been demonstrated to inhibit CRH neuron activity and HPA axis activity.^3^ The mice were stereotaxically injected with 250 nL of Gi DREADD at (−0.9 mm posterior, ±0.3 mm lateral, and 5.1 mm depth), unilaterally injected with kainic acid in the ventral hippocampus (3.6 mm posterior, −2.8 mm lateral, and 2.8 mm depth), and implanted with a prefabricated EEG headmount customized with a ventral hippocampal depth electrode. Electroencephalogram recordings (24/7) began 21 days post-surgery. The first 7 days of EEG recordings were baseline recordings to detect seizure frequency and severity in untreated mice. During the second week of continuous EEG recordings, mice were given *ad libitum* access to 5 mg of clozapine-*N*-oxide (CNO; Sigma-Aldrich cat. #34233-69-7) dissolved in 200 mL of drinking water.

### RU486 administration

After administering vIHKA injections and implanting the prefabricated EEG headmount as described above, a cohort of mice with a 10 mg, 21-day, slow release RU486 pellet (Innovative Research of America, cat. #X-999). Following 3 days of post-surgery recovery, we began continuous, 24/7 EEG recordings.

### Electroencephalogram recordings and analysis

Following vIHKA, we implanted a chronic, custom fabricated EEG headmount (Pinnacle Technology, cat. #8201) outfitted with a coated stainless steel wire depth electrode (A-M Systems, cat. #792300) inserted at the kainic acid injection site, in the right ventral hippocampus. The prefabricated headmount was fixed onto the mouse skull with 3 screws: one serving as an EEG lead in the prefrontal cortex, another as a reference ground, and another as the animal ground. Electroencephalogram recordings were collected at 4 KHz using a 100X gain preamplifier high pass filtered at 1KHz (Pinnacle Technology, cat. #8292-SE) and tethered turnkey system (Pinnacle Technology, cat. #8200). Recordings began between 7-10 days postsurgery and continued uninterrupted for up to 3 weeks. For mice that received Gi DREADD injections, EEG recordings began 21 days following surgery to allow for optimal expression of the Gi DREADDs.

Seizures from EEG traces were automatically detected using a thresholding method, which detected 99.5% of manually scored seizures in a test dataset. Briefly, traces were downsampled (decimated) to 100 Hz and divided into 5 second segments. Seizure segments were detected based on power (2-40 Hz) and line length of the downsampled LFP/EEG signal. These features and their thresholds were selected based on a training dataset along with their performance on the testing dataset (*Feature: feature multiplier*. *Line length*-vHPC: *4.5, Power*-vHPC: *2.5, line length*-FC: *3.5*). 5s segments were scored as seizures if their values were larger than feature mean + (standard deviation x feature multiplier) in at least two out of the three features (popular vote). Seizures were only included if two consecutive 5 second segments were classified as seizures (minimum 10 seconds length). Automatically detected seizures were then manually verified and all false positive events were discarded. These pipelines can now be found in a python application (seizy) that was developed in our laboratory (https://github.com/neurosimata/seizy). Seizure frequency was calculated by dividing the total number of seizures a mouse had over the recording period by the total number of recording hours. The average seizure duration was calculated by taking the mean of seizure bout durations for each mouse in each recording period. For each mouse, seizure burden was calculated as: Total # of seizures * Average seizure durationTotal EEG recording hours

### Behavioral assessments

Behavioral testing was conducted beginning at 60 days post-vIHKA or vehicle injections. Avoidance behaviors were tested using the open field (OF) and light/dark (L/D) paradigms. Learned helplessness behavior was tested using the forced swim test (FST), anhedonia was evaluated using the sucrose preference test (SPT), and motivated, goal-directed behavior was assessed using the nestlet shredding test (NST).

#### Open Field Test

Avoidance behaviors in the light/dark box was measured as previously described (Melon, 2019).^36^ Briefly, mice were placed in an open arena surrounded by a 40 cm x 40 cm photobeam frame with 16 x 16 equally spaced photocells (Hamilton-Kinder). Mice were placed individually into the center of the arena and beam breaks and movement were automatically detected and measured using the Motor Monitor software (Hamilton-Kinder) over the 10 min testing period. We analyzed the amount of time spent in the center, the total distance traveled in the center, and the total number of beam breaks.

#### Light/Dark Box

Avoidance behaviors in the light/dark box was measured as previously described (Melon, 2019).^36^ Briefly, mice were placed in the dark chamber of a two-chamber light/dark box apparatus surrounded by a 22 cm x 43 cm photobeam frame with 8 equally spaced photocells (Hamilton-Kinder). Mice were allowed to explore the two-chamber apparatus freely for a 10 min testing period. Beam breaks and movement were detected and measured using the automated Motor Monitor software (Hamilton-Kinder). We analyzed the total number of entries, total duration spent in, and the total distance traveled in the light chamber.

#### Sucrose Preference Test

Anhedonia was measured using the sucrose preference test, using a protocol similar to our previous studies (Melon, 2019).^36^ Mice were individually housed and given *ad libitum* access to two water bottles, one filled with water and the other filled with 2% sucrose (w/v). Water bottles were weighed daily at the same time (ZT 14-15). Water bottle positions were swapped daily to avoid placement preference. The mice underwent the SPT paradigm for 7 days. Sucrose preference for each mouse was calculated by dividing the daily change in the sucrose water bottle weight by the sum of the changes in the bottle weights for both the regular water and sucrose water (Δ sucrose/Δ sucrose + Δ reg. water). Graphs were generated by plotting the average sucrose preference across the 7 days for each mouse. (1 gram of H_2_0 = 1 mL of H_2_0).

#### Nestlet Shredding Test

Mice were placed in clean cages with two squares of unshredded cotton fiber nestlets (Ancare catalog #NES3600). Before the mice were placed in the cage, the unshredded nestlet was weighed. Any unshredded nestlet was weighed at the same time every day (ZT 14-15) for 7 days. Care was taken to not disrupt the unshredded nestlet; any piece that was physically attached to the nestlet counted towards the overall weight of the nestlet. The percent of the nestlet left unshredded was calculated by taking the weight of any unshredded nestlet for each day and dividing it by the initial weight of the nestlet (Day 0).

#### Forced Swim Test

Learned helplessness was measured using the FST as previously described by our laboratory.^36,37^.Briefly, mice were placed in a 5 L circular, plastic container (20 cm in diameter). The container was filled with room temperature water (23-25° C) and placed in a room with minimal visual and auditory distractions. Mice were individually placed in water and were video recorded for 6 minutes, undisturbed. FST videos were analyzed by a FST Scoring application developed in our laboratory. The latency to the first bout of immobility and total time spent immobile was quantified.

#### Automated Behavioral Testing

Mouse tracking for FST was done using a custom trained DeepLabCut model.^38^ Mouse nose, tail, tail-base and rear feet were automatically labeled, and positional data was processed to exclude and forward-fill low confidence values. Distance traveled between frames for each limb was used to determine if the mouse was immobile based on a machine-learning model. The model used a Long Short-Term Memory network (https://github.com/fchollet/keras) which was trained using manually scored experiments from several lab members. After excluding bouts of immobility under 1.5 seconds, total time immobile and latency to first bout of immobility were calculated. Values from experiments manually verified *post hoc* correlated with the automated analysis with an R^2^ of 0.92 (n=9). Python scripts used to process and analyze the data, as well as the artificial intelligence models used are available upon request.

#### Behavioral distribution analysis by peak detection

Kernel Density Estimations (KDE) were fit to behavioral data using the Seaborn package for Python (https://github.com/mwaskom/seaborn). The default parameters were used, except the bandwidth was changed to 0.8 to adjust for over-smoothing of the distributions. Peaks were then detected using the Scipy *Find Peaks* function. The midpoint between the peaks was used as the threshold to determine if each animal fell into the vulnerable or non-vulnerable populations. If only one peak was detected, the mean of the values was used to split the animals into vulnerable and non-vulnerable groups.

### Corticosterone ELISAs

Blood was collected from mice either through submandibular bleeds using a mouse lancet^39^ or through trunk blood at the end of the experimental timeline. Blood collection was performed at the same time (ZT 10-11). During collection, samples were kept on ice. Blood samples were then centrifuged at 1.8 x *g* for 15 minutes at 4°C to isolate serum. Serum was stored in the −80°C freezer until use. Postmortem blood samples from PWE, PWE with suspected SUDEP, and subjects with no history of epilepsy were obtained from the North American SUDEP Registry (NASR) at NYU Langone Health. To quantify corticosterone and corticotropin releasing hormone (CRH) levels from our serum samples, we used a corticosterone enzyme linked immunosorbent assay (ELISA) kit (Enzo Life Sciences, cat. #ADI-900-097) or a Human CRH/CRH ELISA kit (Lifespan Biosciences, cat. #LS-F5352) and ran 5 uL aliquots of each sample in duplicate. Experimental samples were compared to a standard curve of known corticosterone concentration.

### Immunohistochemistry

Following the completion of EEG recordings or behavior, mice were sacrificed by anesthetizing with isoflurane and euthanized by rapid decapitation. The brain was rapidly extracted, immersed in 4% paraformaldehyde, and incubated at 4°C overnight. Following immersion fixation, the brains were cryoprotected in 10% and 30% sucrose at 4°C for 24 and 48 hours, respectively, snap frozen, and stored at −80° C until cryostat sectioning.

Brains were coronally sectioned on a crysostat at 40 μm. For NeuN staining, sections were blocked for 1 hour in 10% normal goat serum (at room temperature) and probed with an anti-NeuN Alexa Fluor 488 conjugated antibody (1:100; Millipore-Sigma cat. # MAB377X) for 2 hours at room temperature and protected from light. For ZnT3 staining, endogenous peroxidase activity in the tissue was quenched by incubating sections in 3% hydrogen peroxide in methanol for 30 minutes. Sections were then blocked in 10% normal goat serum for one hour at room temperature. Sections were incubated in primary anti-ZnT3 (Synaptic Systems, cat. # 197002; 1:5000) for 72 hours at 4°C. Goat-anti Rabbit Vectastain Elite ABC-HRP kit (Vector Laboratories, cat.# PK-6101) was used according to manufacturer’s instructions and 3,3’ Diaminobenzidine (DAB) substrate was used for visualization.

The NeuN staining was visualized using a Nikon A1R confocal. Each dentate gyrus image was imaged at 20X and stitched together using the Fiji software.^40^ Brightfield images of the ZnT3 staining were taken on a Nikon E800 microscope. Three hippocampal sections (representing the dorsal, medial, and ventral sections of the hippocampus) were imaged and analyzed per mouse. Each hippocampal section was then normalized to its respective uninjected, contralateral side. The average per mouse was then used to average across animals.

#### Dentate granule cell dispersion and Mossy Fiber Sprouting quantification

Cell dispersion was quantified using Cell Profiler ^41^ to detect individual cells expressing NeuN using the included cell profiler pipelines. Briefly, the Otsu detection method with global, three-class thresholding was used to detect cells from the DAPI channel. The threshold correction factor was adjusted between 1.3 and 2 to manually adjust for variance in tissue thickness. A NeuN mask was then used to select only neuronal cells. Cell density was calculated by dividing the total number of cells by the area of manually outlined dentate gyrus. Dentate granule cell dispersion was analyzed by quantifying the average number of neighboring cells and the total percent of cells each dentate granule cell was touching in an unbiased fashion.

Mossy fiber sprouting (MFS) was also quantified using an included Cell Profiler pipeline.^41^ Briefly, the Otsu detection method with global, two-class thresholding was used to identify ZnT3 stained mossy fibers. MFS was quantified as the measure of the total percent of mossy fiber sprouting in the dentate gyrus in an unbiased manner. Total mossy fiber area was divided by the area of the manually outlined dentate gyrus.

### Statistical analyses

Data was analyzed using GraphPad Prism 9. When only two conditions were present, we ran a Student’s *t*-test. Multi-group analyses were tested with a one-way ANOVA with post-hoc multiple comparisons. When comparing multiple conditions and treatments, a two-way ANOVA test with post-hoc Tukey’s multiple comparisons was used. All data are represented as the mean ± SEM. All *p*-values <0.05 were considered significant. \**p* < 0.05; \*\**p* < 0.01; \*\*\**p* < 0.001; \*\*\*\**p* < 0.0001

## Results

To examine the impact of HPA axis hyperexcitability on epilepsy outcomes, we used a mouse model that exhibits exacerbated seizure-induced activation of the HPA axis (*Kcc2/Crh* mice) and evaluated the impact on several epilepsy outcome measures, including spontaneous recurrent seizure frequency, characteristic neuropathological features, and psychiatric comorbidities.

### Seizure-induced activation of the HPA axis in *Kcc2/Crh* mice

The *Kcc2/Crh* mouse model was previously generated and characterized by our laboratory where we demonstrated that these animals have increased stress-induced elevations in corticosterone levels compared to controls Cre^-/-^ littermates (WT).^36^ Here we demonstrate that *Kcc2/Crh* mice also exhibit exacerbated seizure-induced elevations in corticosterone and elevated circulating corticosterone levels in chronically epileptic mice (Fig. 1A). Corticosterone levels are increased in *Kcc2/Crh* mice two hours following acute kainic acid (20 mg/kg) administration (237.52 ± 39.66 ng/ml) compared to WT controls (100.48 ± 20.148 ng/ml) and vehicle controls (49.01 ± 8.58 ng/ml) (Fig. 1A). Circulating corticosterone levels are also increased in *Kcc2/Crh* mice following *status epilepticus* induction using the pilocarpine model (Fig. 1A; 259.8 ± 79.69 ng/ml) compared to chronically epileptic WT mice (Fig 1B; 71.07 ± 10.85). Thus, this mouse model (*Kcc2/Crh*) is a valuable tool for examining the pathophysiological impact of HPA axis hyperexcitability on epilepsy outcomes.

**Figure 1:**
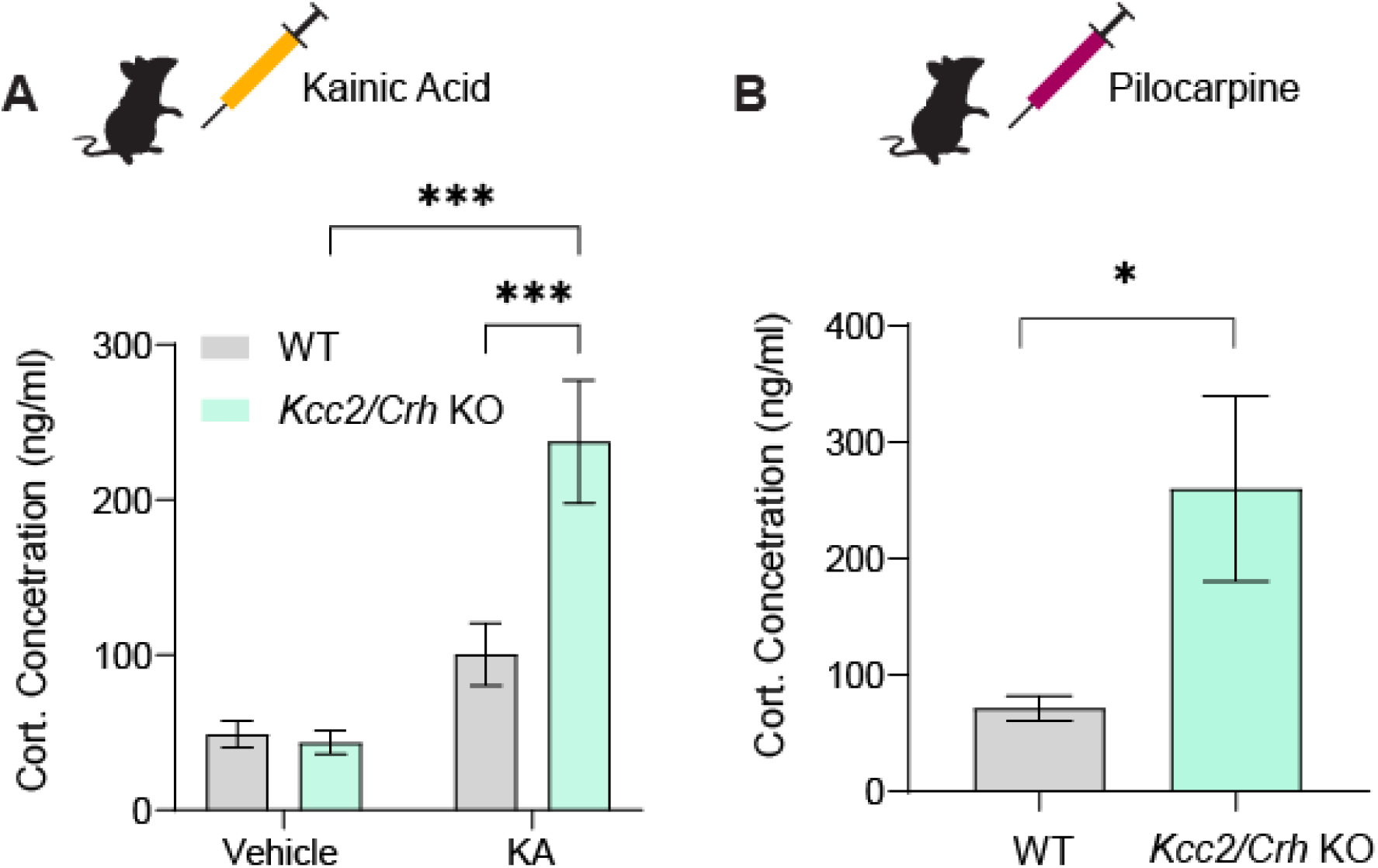
*Kcc2/Crh* KO mice exhibit an exaggerated, seizure-induced activation of the HPA axis. **(A)** Circulating corticosterone concentration collected 2 hours after intraperitoneal injections of either vehicle or KA in WT and *Kcc2/Crh* mice. **(B)** Circulating corticosterone levels were quantified from serum collected 24 hours following pilocarpine induced *status epilepticus* in WT and *Kcc2/Crh* mice. n = 20 (WT vehicle); 5 (*Kcc2/Crh* vehicle); 19 (WT KA); 8 (*Kcc2/Crh* KA); 12 (WT pilocarpine); 9 (*Kcc2/Crh* pilocarpine). Error bars represent ±SEM. \**p* <0.05; \*\*\**p*<0.001. A two-way ANOVA was used to analyze **(A)** and an unpaired Student’s *t*-test was used to analyze **(B)**. WT, wildtype; KA, kainic acid; Cort., corticosterone.

### HPA dysfunction worsens mossy fiber sprouting in chronically epileptic *Kcc2/Crh* mice

Hallmark neuropathological features of temporal lobe epilepsy (TLE), including hippocampal mossy fiber sprouting (MFS) and dentate granule cell dispersion (DGCD), have been well characterized in both preclinical models of epilepsy and in PWE.^42–46^ Chronic pathological activation of the HPA axis in clinical models of mood disorders has also been shown to compromise hippocampal function and integrity.^47–51^ Here, we assessed whether HPA axis dysfunction additively worsened characteristic neuropathology in chronically epileptic *Kcc2/Crh* mice. Eight weeks following either vIHKA or vIHSa, brains from WT and *Kcc2/Crh* mice were collected, processed, and stained with either ZnT3 (for MFS) or DAPI and NeuN (for DGCD). Compared to chronically epileptic WT mice, a significantly greater percent of the dentate gyrus subregion of the hippocampus in vIHKA *Kcc2/Crh* mice contains MFS (Fig. 2A and D; WT vIHKA: n= 11, 2.546 ± 0.4240%; *Kcc2/Crh* vIHKA: n= 9, 11.85 ± 5.367%; unpaired Student’s *t*-test, \**p*< 0.05).

**Figure 2:**
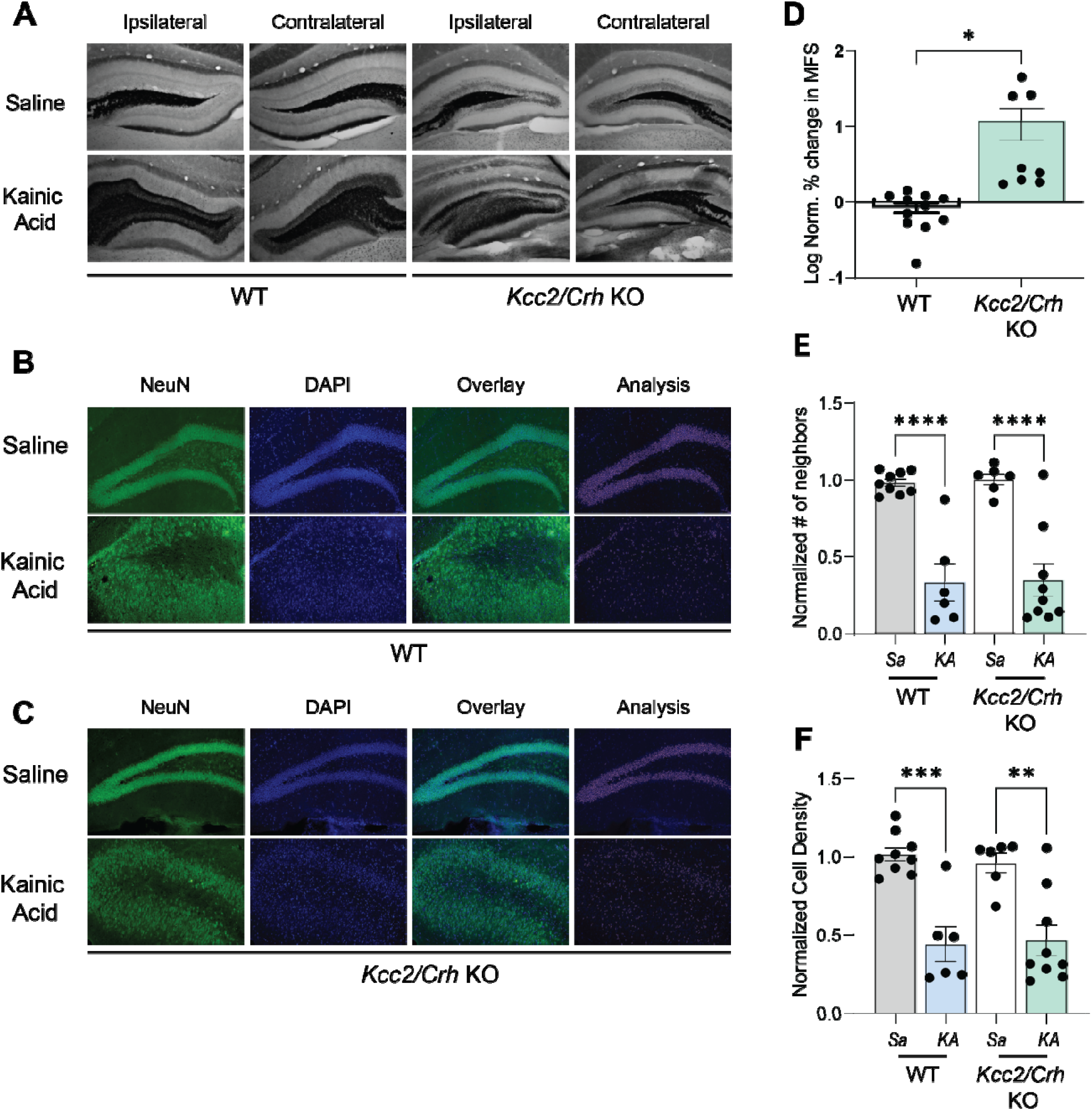
HPA axis dysfunction worsens MFS in chronic epilepsy. **(A)** Representative coronal sections of the hippocampus collected from control and chronically epileptic adult, male WT and *Kcc2/Crh* KO mice and stained with ZnT3 to quantify MFS. **(B-C)** Representative sections stained with NeuN to visualize DGCD in WT and *Kcc2/Crh* mice are shown in **(B)** and **(C)**, respectively. Pink outlines were automatically generated through Cell Profiler and indicate cells where NeuN and DAPI colocalize. **(D)** The mean (±SEM) percent change in MFS was quantified in the ipsilateral hemisphere and normalized to the mean percent change in MFS of the contralateral hemisphere of chronically epileptic WT and *Kcc2/Crh* mice. **(E)** The mean (±SEM) number of adjoining neighboring neuronal cells was quantified for the ipsilateral hemisphere and normalized to the mean number of immediate neighboring neuronal cells on the contralateral hemisphere for both control and chronically epileptic WT and *Kcc2/Crh* mice. **(F)** The total number of cells within the manually defined dentate gyrus area was quantified on the ipsilateral hippocampal hemisphere and normalized to the cell density of the uninjected, contralateral hippocampal hemisphere. *N*= 2-3 mice per group; *n*= 6-9 sections per group, with 3 regionally distinct hippocampal sections from each mouse (dorsal, medial, and ventral). Error bars represent ±SEM. An unpaired Student’s *t*-test was used to analyze MFS and a one-way ANOVA was used to analyze DGCD parameters. Statistical analysis was performed on the sample size, *n*, for both MFS and DGCD. \**p* < 0.05; \*\**p* < 0.01; \*\*\**p* < 0.001; \*\*\*\**p* < 0.0001. WT, wild type; KA, kainic acid; MFS, mossy fiber sprouting; DGCD, dentate granule cell dispersion; Norm, normalized.

To quantify the extent of DGCD in chronically epileptic WT and *Kcc2/Crh* mice, we analyzed the mean number of cells in direct contact with cells co-expressing DAPI and NeuN in addition to calculating the average cell density. While there were significant differences in the mean number of immediately adjacent neighbors between the vIHSA and vIHKA injected WT and *Kcc2/Crh* groups (Fig. 2B, C, and E; WT vIHSA: n= 9, 0.982 ± 0.024; WT vIHKA, n= 6, 0.332 ± 0.122; *Kcc2/Crh* vIHSa: n= 6, 1.005 ± 0.037; *Kcc2/Crh* vIHKA, n= 9, 0.345 ± 0.1071 normalized number of neighbors; \*\*\*\**p* < 0.0001), there were no statistical differences in the mean number of adjacent neighbors between the WT and *Kcc2/Crh* vIHKA groups (Fig. 2B, C, and E). Similarly, there were significant differences in the total cell density of the dentate granule cell region between the vIHSa and vIHKA injected WT and *Kcc2/Crh* groups (Fig. 2B, C, and F; WT vIHSA: n= 9, 1.016 ± 0.044; WT vIHKA, n= 6, 0.441 ± 0.112; *Kcc2/Crh* vIHSa: n= 6, 0.960 ± 0.063; *Kcc2/Crh* vIHKA, n= 9, 0.466 ± 0.099 normalized cell density; \*\**p*< 0.01, \*\*\**p*< 0.001), there were no statistical differences between the WT and *Kcc2/Crh* vIHKA groups (Fig. 2B, C, and F). Because HPA axis function is already aberrantly increased in chronic epilepsy, further exacerbating HPA axis dysfunction in the *Kcc2/Crh* mice appears to selectively worsen MFS, but not DGCD, in chronic epilepsy.

### Chronically epileptic *Kcc2/Crh* mice exhibit increased vulnerability to negative affective states

Previous studies have demonstrated behavioral abnormalities, including increased avoidance behaviors and anhedonia, in the ventral intrahippocampal kainic acid (vIHKA) model of chronic epilepsy.^35^ Therefore, we employed this model to assess the impact of HPA axis dysfunction on behavioral abnormalities associated with chronic epilepsy. Eight weeks after vIHKA or vIHSa injections, chronically epileptic and control *Kcc2/Crh* and Cre^-/-^ (WT) mice were subjected to a battery of behavioral paradigms to test for differences in behavioral states. Avoidance behaviors were tested in the open field (OF) and light/dark (L/D) box paradigms. Compared to vIHSa mice, chronically epileptic *Kcc2/Crh* and WT mice show statistically significant decreases in the amount of time spent (WT vIHSa: n =17, 99.34±14.47 s; *Kcc2/Crh* vIHSa: n =20, 65.15 ± 8.623 s; WT vIHKA: n= 11, 50.25 ± 8.496 s; *Kcc2/Crh* vIHKA: n=19, 45.59 ± 7.416 s; one-way ANOVA; \**p* < 0.05, \*\*\**p* < 0.001) and total distance traveled (WT vIHSa: n=17, 1009 ± 107.5 cm; *Kcc2/Crh* vIHSa: n=20, 584.1 ± 69.48 cm; WT vIHKA: n=11, 647.7 ± 110.0 cm; *Kcc2/Crh* vIHKA: n=19, 509.8 ± 69.98 cm; one-way ANOVA; \**p* < 0.05, \*\**p* <0.01, \*\*\**p* < 0.001) in the center of the OF test (Fig. 3A and B), both of which are indications of increased avoidance behaviors. While there are no statistically significant differences in the amount of time that was spent in the light arena of the L/D test across the groups (Fig. 3C; WT vIHSa: n=17, 159.2 ± 14.14 s; *Kcc2/Crh* vIHSa: n=20, 131.2 ± 17.55 s; WT vIHKA: n=11, 136.0 ± 22.99 s; *Kcc2/Crh* vIHKA: n=19, 105.9 ± 17.81 s; not significant via one-way ANOVA), there was a modest, but statistically significant, decrease in the total distance the chronically epileptic *Kcc2/Crh* mice traveled in the light compared to the saline injected WT mice (Fig. 3D; WT vIHSa: n=17, 1073 ± 84.79 cm; *Kcc2/Crh* vIHSa: 790.5 ± 92.88 cm; WT vIHKA: 924.0 ± 173.9 cm; *Kcc2/Crh* vIHKA: 675.2 ± 111.3 cm; one-way ANOVA; \**p* < 0.05). These modest behavioral effects observed in *Kcc2/Crh* mice may represent a potential floor effect in the degree of behavioral severity that chronically epileptic mice will exhibit, since the HPA axis is elevated in chronically epileptic mice as well and a further increase may not translate into a further impact on behavior.

**Figure 3:**
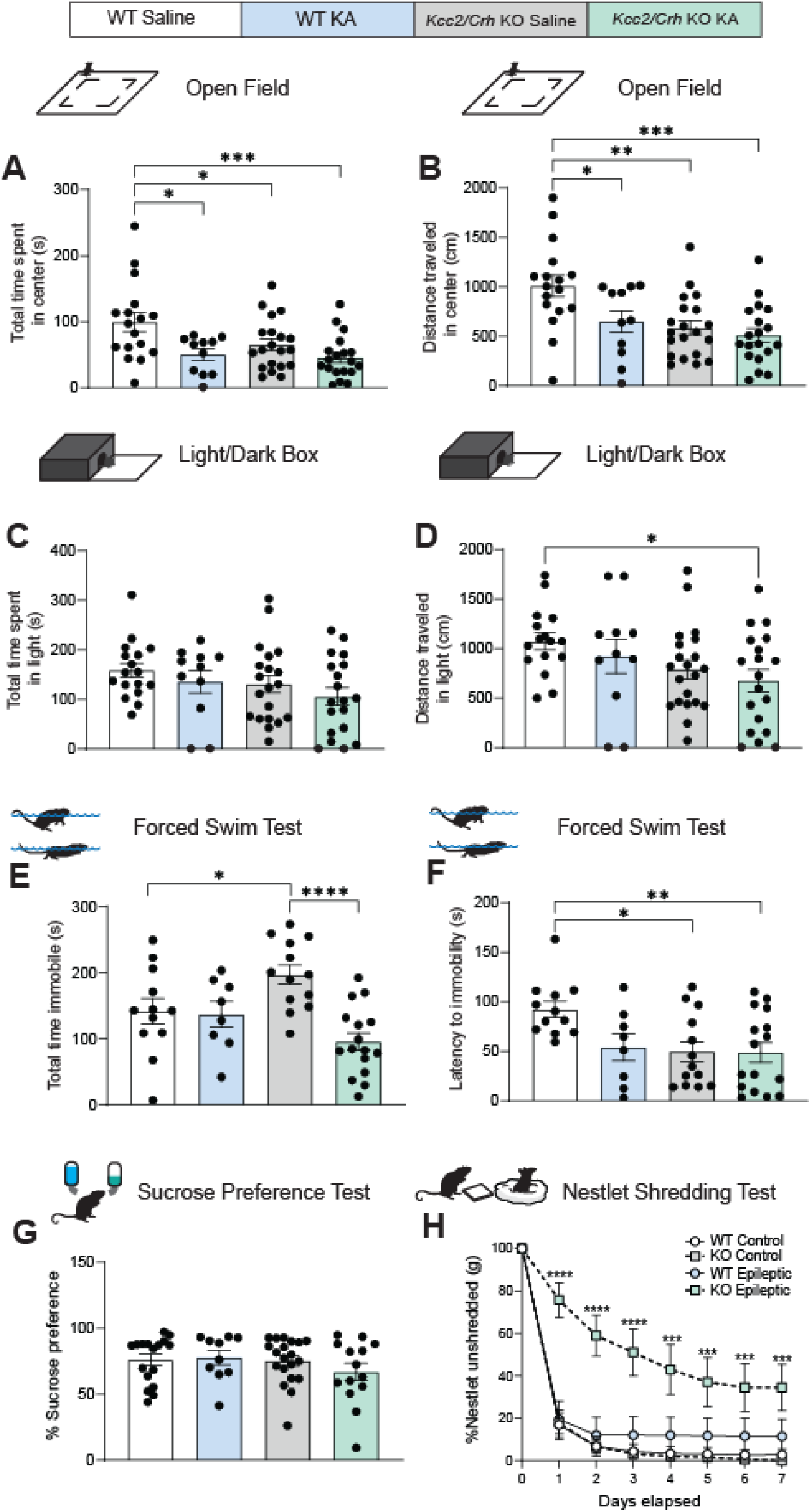
Chronically epileptic *Kcc2/Crh* male mice exhibit increased vulnerability to negative affective states compared to WT control mice. **(A-B)** The average total time spent in the center **(A)** and total distance traveled in the center of the OF test **(B)**. **(C-D)** The average total time spent **(C)** and total distance traveled **(D)** in the lit arena of the LD setup. **(E-F)** The histograms depict the average total time spent immobile **(E)** and the latency to the first bout of immobility **(F)** in the FST. **(G)** The average percent sucrose preference measured over the course of a 7-day SPT paradigm. **(H)** The average percent of unshredded nestlet material was weighed daily for one week. *N*= 11-20 per group. Error bars represent SEM. \**p* < 0.05; \*\**p* < 0.01; \*\*\**p* <0.001; \*\*\*\**p* <0.0001. OF, LD, FST, and SPT were analyzed using a one-way ANOVA. NST was analyzed using a two-way ANOVA with repeated measures. WT, wildtype; KA, kainic acid; OF, open field; LD, light/dark; FST, forced swim test; SPT, sucrose preference test; NST, nestlet shredding test.

**Figure 4:**
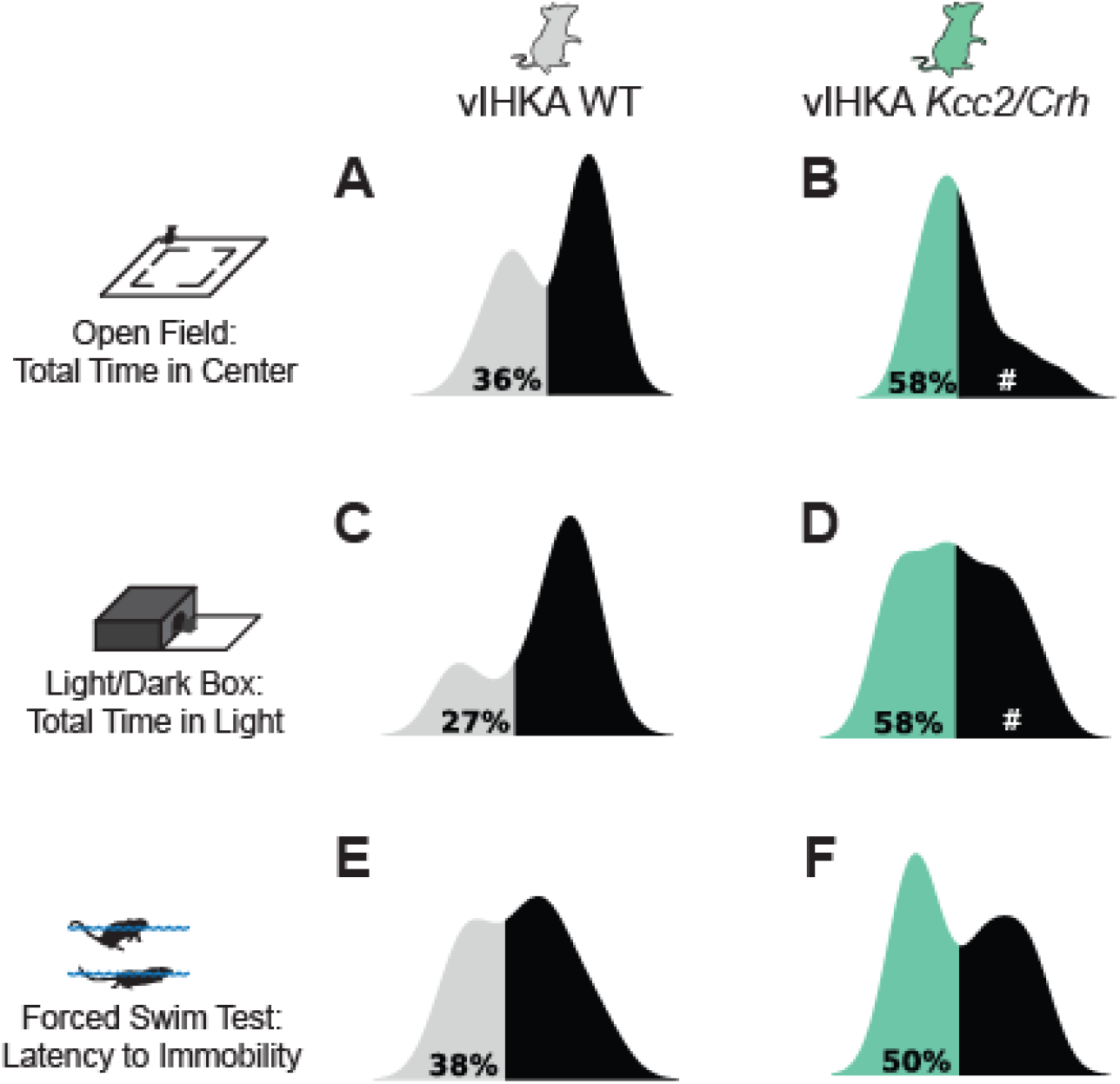
A greater proportion of the chronically epileptic *Kcc2/Crh* mouse population exhibit increased vulnerability to negative affective states compared to chronically epileptic WT mice. **(A-B)** Population distributions of performance in the OF test between chronically epileptic WT **(A)** and *Kcc2/Crh* **(B)** groups. **(C-D)** Distribution of performance in the LD box between the vIHKA WT **(C)** and vIHKA *Kcc2/Crh* **(D)** groups. **(E-F)** Within group distributions of performance in the FST test between the chronically epileptic WT **(E)** and *Kcc2/Crh* **(F)** groups. In each plot, the lighter color represents the underperforming, more vulnerable population while the black distribution plots represent the resilient groups. # denotes instances where only one peak was detected, so population distributions were delineated by the mean of the data. OF, open field; LD, light dark box; FST, forced swim test.

Because hyperactivation of the HPA axis is a cardinal feature in major depression models, we hypothesized that the exaggerated seizure induced activation of the HPA axis in the *Kcc2/Crh* mice would correlate with greater susceptibility for behavioral outcomes potentially relevant to depression, including learned helplessness, anhedonia, and decreased motivation for goal-directed behaviors, associated with chronic epilepsy. Behavioral outcomes were evaluated in control and chronically epileptic WT and *Kcc2/Crh* mice using a battery of behavioral approaches including the forced swim test (FST; a test of learned helplessness), ^52^ sucrose preference test (SPT; a test of anhedonia),^53^ and nestlet shredding test (NST; a measure of apathy or loss of goal directed behavior).^54^ *Kcc2/Crh* mice exhibit an increase in total time spent immobile in the forced swim test compared to saline-injected WT mice, an effect which is lost in chronically epileptic mice (Fig, 3E; WT vIHSa: n=12, 142.0 ± 19.29 s; *Kcc2/Crh* vIHSa: n=13, 197.4 ± 14.33 s; WT vIHKA: n=8, 137.1 ± 19.43 s; *Kcc2/Crh* vIHKA: n=16, 95.71 ± 13.03 s; one-way ANOVA; \**p* < 0.05, \*\*\*\**p* < 0.0001). Chronically epileptic *Kcc2/Crh* mice exhibit a significant decrease in the latency to immobility in the FST compared to saline injected WT mice (Fig. 3F; WT vIHSa: n=12, 92.37 ± 8.194 s; *Kcc2/Crh* vIHSa: n= 13, 49.52 ± 10.32 s; WT vIHKA: n=8 53.96 ± 13.71 s; *Kcc2/Crh* vIHKA: n=16 48.79 ± 9.801 s; one-way ANOVA; \**p* < 0.05, \*\**p*< 0.01). The increase in learned helplessness behavior within the chronically epileptic *Kcc2/Crh* mice is mirrored in the chronically epileptic WT group, once again suggesting that in chronic epilepsy, there is a floor effect in which behavioral severity is linked to activation of the HPA axis but is independent of the magnitude of HPA axis activation. Similarly, there were no observable differences in anhedonic behavior as measured by the sucrose preference test in either the control or chronically epileptic *Kcc2/Crh* and WT mice.

Chronically epileptic *Kcc2/Crh* mice also exhibit profound deficits in motivated goal-directed behaviors, evident from the NST (Fig. 3H). Shredding nestlet or bedding to build a nest is an innate, goal directed behavior in rodents.^54^ When presented with an intact nestlet, chronically epileptic *Kcc2/Crh* mice lag in shredding their nestlet over the course of one week compared to vIHSa mice and vIHKA WT mice (Fig. 3H). A two-way ANOVA with repeated measures analysis reveals a significant interaction effect of KA treatment and the percent of unshredded nestlet remaining over time (Fig. 3H; \*\*\**p*< 0.001, \*\*\*\**p* <0.0001), an effect that is solely driven by the chronically epileptic *Kcc2/Crh* group. In summary, chronically epileptic *Kcc2/Crh* mice show signs of greater susceptibility to apathy and learned helplessness, both of which are symptoms of depression-like behavior, compared to chronically epileptic WT mice.

Consistent with the role of HPA axis dysfunction driving behavioral deficits in chronically epileptic mice, the vIHSa *Kcc2/Crh* mice, which exhibit HPA axis dysfunction but are not chronically epileptic, also show behavioral abnormalities. Compared to vIHSa WT mice, vIHSa *Kcc2/Crh* mice exhibit significant decreases in the total time spent (Fig. 3A; WT vIHSa: n =17, 99.34±14.47 s; *Kcc2/Crh* vIHSa: n =20, 65.15 ± 8.623 s; one-way ANOVA; *p* < 0.05) and in the total distance traveled (Fig. 3B; WT vIHSa: n=17, 1009 ± 107.5 cm; *Kcc2/Crh* vIHSa: n=20, 584.1 ± 69.48 cm; one-way ANOVA; *p* < 0.05) in the center during the OF paradigm. Additionally, vIHSA *Kcc2/Crh* mice spend a significantly greater time immobile (Fig. 3E; WT vIHSa: n=12, 142.0 ± 19.29 s; *Kcc2/Crh* vIHSa: n=13; one-way ANOVA; *p* < 0.05) and exhibit faster latency to immobility (Fig. 3F; WT vIHSa: n=12, 92.37 ± 8.194 s; *Kcc2/Crh* vIHSa: n= 13, 49.52 ± 10.32 s; one-way ANOVA; *p* < 0.05) in the FST compared to vIHSA WT mice. These data suggest that overactivation of the HPA axis alone is sufficient to alter negative affective states.

It is also interesting to note that there is substantial variance in the data within each group across these behavioral paradigms. This prompted us to hypothesize that there may be two populations within each group, one that is vulnerable and another that is resilient to altered affective states associated with epilepsy. While none of the distributions passed a Hartigan-Dip test for bimodality, likely due to the data being underpowered, we plotted the distribution of the populations for the chronically epileptic *Kcc2/Crh* and WT by using a kernel density estimate (KDE) to determine a continuous probability density curve. From the relative probability distribution that was generated using the KDE, peaks of the population were automatically detected in an unbiased manner, and the population was delineated by these peaks into either a vulnerable (those that underperformed in a particular behavioral task compared to vIHSa WT mice) or resilient group. If the algorithm did not detect peaks, the population was split based on the mean of the distribution. Using this algorithm, we found that compared to chronically epileptic WT mice, a greater proportion of the chronically epileptic *Kcc2/Crh* mice were vulnerable across the behavioral paradigms (Fig.4). For example, when looking at the total time the chronically epileptic mice spent in the center of the OF test, the algorithm we used indicates that 58% of the *Kcc2/Crh* mice are considered vulnerable compared to just 36% of the WT group (Fig.4A and B). Similarly, 58% of the chronically epileptic *Kcc2/Crh* mice are considered more vulnerable to anxiety-like behavior in the total time spent in the light arena of the L/D box compared to just 27% of the chronically epileptic WT mice (Fig.4C and D). In summary, our data demonstrates that HPA axis dysfunction in the *Kcc2/Crh* mice increases vulnerability to negative affective states associated with chronic epilepsy; however, these effects are modest and likely due to a floor effect.

### Chronically epileptic *Kcc2/Crh* mice have increased SUDEP incidence

Given the proconvulsant actions of stress hormones,^23,26,27^ we sought to determine whether seizure induced activation of the HPA axis also leads to the worsening of seizure outcomes. At the time of vIHKA administration, WT and *Kcc2/Crh* mice were implanted with EEG head mounts outfitted with a ventral hippocampal depth electrode. Spontaneous seizures were observed 7 days following vIHKA-induced *status epilepticus*. Between chronically epileptic WT and *Kcc2/Crh* mice, there were no significant differences in the number of daily seizures (Fig. 5C; WT vIHKA: N= 6; 1.924 ± 0.542 seizures; *Kcc2/Crh* vIHKA: N= 8; 2.885 ± 0.848 seizures; no significance via one-way ANOVA), average seizure duration (Fig. 5D; WT vIHKA: N= 7; 54.20 ± 3.97 s; *Kcc2/Crh* vIHKA: N= 8; 53.49 ± 2.213 s; no significance via one-way ANOVA), or seizure burden (Fig. 5E; WT vIHKA: N= 6; 4.267 ± 1.15; *Kcc2/Crh* vIHKA: N =8; 6.539 ± 1.984). Although the impact of HPA axis hyperexcitability has no significant impact on spontaneous seizure activity, remarkably, we found that 38% of *Kcc2/Crh* mice died of sudden unexpected death in epilepsy (SUDEP) within 20 days post SE (Fig. 6A; vIHKA *Kcc2/Crh*: N= 13; 15.80 ± 2.746 days). In the same time frame, there were no deaths in the chronically epileptic WT mouse population (Fig. 6A; WT vIHKA: N=7). This is an unexpected, yet transformative discovery, potentially linking HPA axis dysfunction to SUDEP risk.

**Figure 5:**
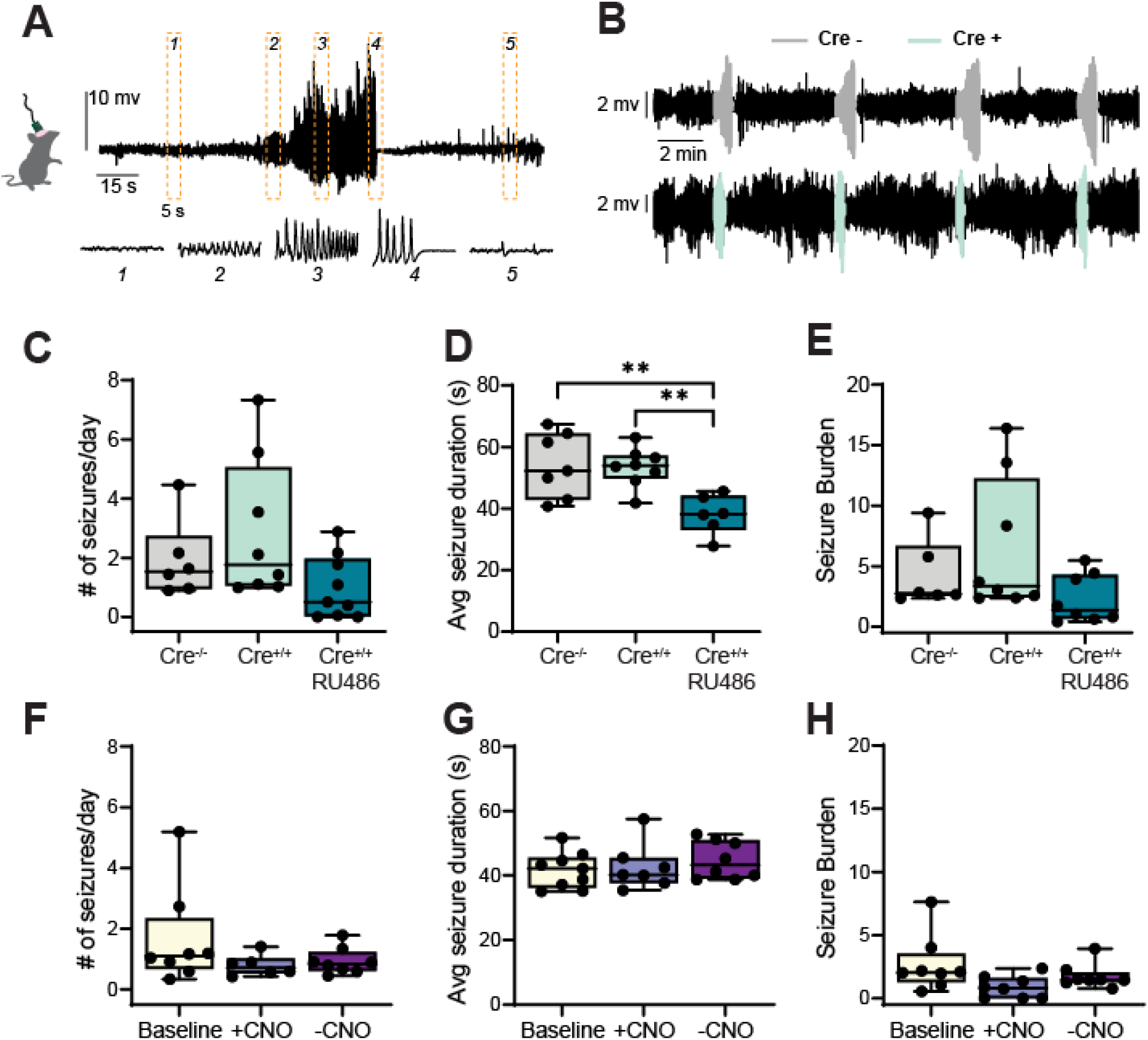
HPA axis dysfunction does not worsen spontaneous seizure activity in chronic epilepsy. **(A)** Representative recorded seizure from hippocampal local field potential. Orange squares indicate time magnified 5s traces. **(B)** Example detected hippocampal seizures which were concatenated from a WT (Cre^-/-^) and a *Kcc2/Crh* KO (Cre^+/+^) mouse. **(C-E)** Mean (±SEM) number of daily seizure occurrences **(C)**, seizure duration **(D)**, and seizure burden **(E)** for chronically epileptic WT, *Kcc2/Crh* KO, and *Kcc2/Crh* KO mice treated with RU486, a glucocorticoid receptor antagonist. **(F-H)** Average number of daily seizures **(F)**, seizure durations **(G)** and seizure burden **(H)** for vIHKA *Kcc2/Crh* mice bilaterally injected with hM4D(Gi)-DREADDs in the PVN. Graphs **F-H** depict mean seizure activity collected via EEG recordings that were made prior to CNO administration (baseline), during CNO administration (+CNO), and during a week free of CNO administration (-CNO). Seizure burden was calculated by multiplying the total number of seizures a mouse exhibited by their average seizure duration, and dividing this value by the total EEG recording hours. Error bars represent SEM. \*\**p* < 0.01. Data was analyzed through a one-way ANOVA. N=5-9 per group. CNO, clozapine-N-oxide; Avg, average.

**Figure 6:**
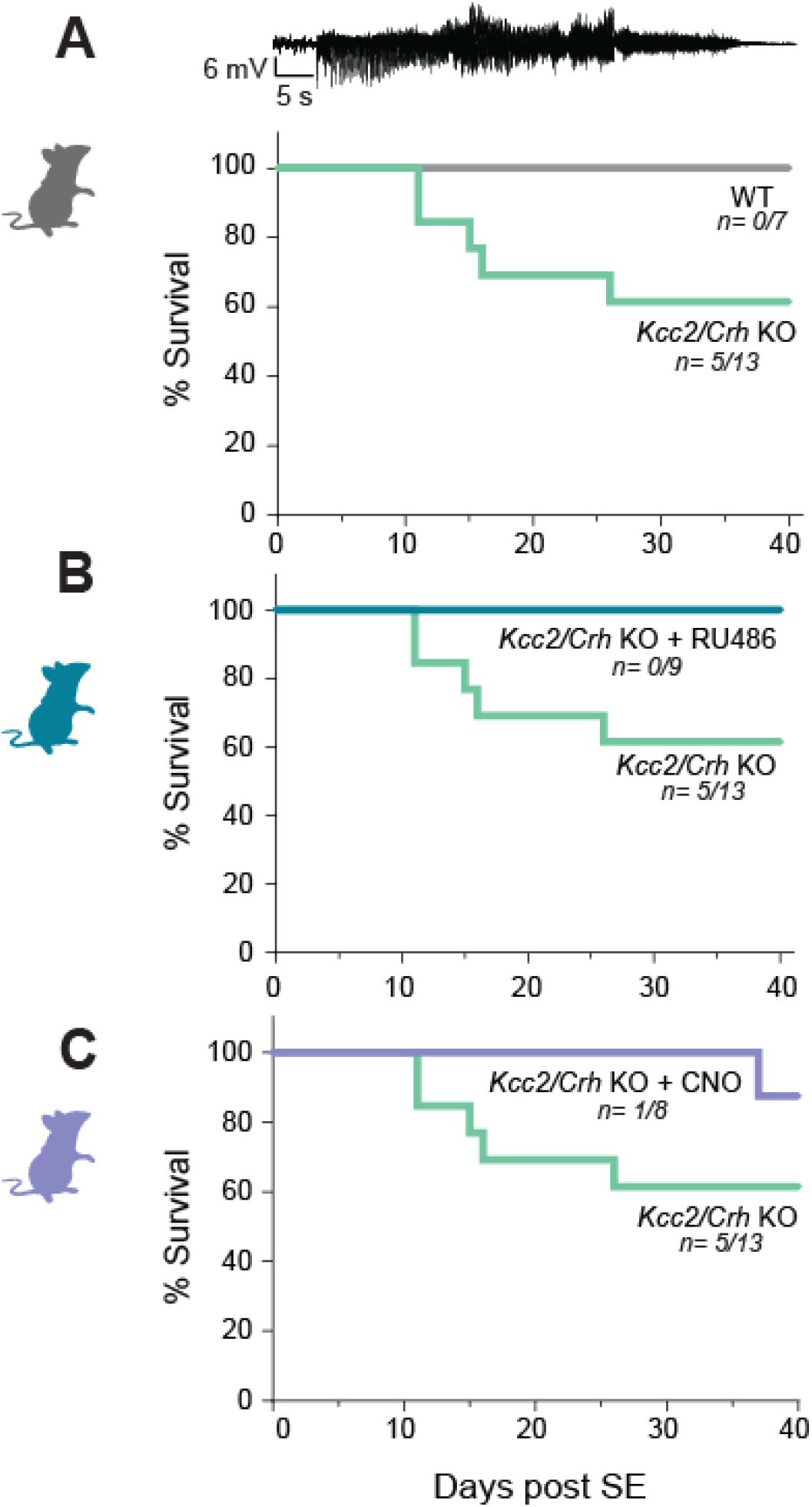
HPA axis dysfunction in chronically epileptic *Kcc2/Crh* mice increases SUDEP incidence. **(A-C)** Percent survival following vIHKA induced SE in chronically epileptic WT and *Kcc2/Crh* KO mice **(A)**, in chronically epileptic *Kcc2/Crh* KO mice treated with the glucocorticoid receptor antagonist, RU486 **(B)**, and in vIHKA *Kcc2/Crh* KO mice expressing Gi DREADDs and given CNO **(C)**. SUDEP, sudden unexpected death in epilepsy; WT, wildtype; SE, status epilepticus.

To further interrogate whether increased HPA axis dysfunction in the *Kcc2/Crh* mice contributes to the increased risk of SUDEP in *Kcc2/Crh* mice, we used chemogenetic and pharmacologic approaches to attenuate HPA axis activity. Our first approach was to stereotaxically inject hM4D(Gi) coupled inhibitory designer receptors exclusively activated by designer drugs (Gi DREADDs) into the paraventricular nucleus (PVN) of the hypothalamus in *Kcc2/Crh* mice immediately prior to the vIHKA injection, an approach previously demonstrated to suppress CRH neuronal activity in the PVN^3^ and decrease seizure frequency.^24^ Three weeks after SE, we collected baseline EEG activity from chronically epileptic *Kcc2/Crh* mice. The following week, the Gi injected *Kcc2/Crh* mice were given the synthetic ligand, clozapine-N-oxide (CNO), via drinking water to inhibit CRH neuron activity. Though inhibition of CRH neuronal activity did not have any impact on overall seizure frequency, duration, or burden in this mouse model of chronic epilepsy (Fig. 5F-H; vIHKA *Kcc2/Crh* + CNO: N= 8; seizures per day: 1.314 ± 0.6537; seizure duration: 42.69 ± 2.753; seizure burden: 1.753 ± 0.341) compared to baseline (Fig. 5F-H; vIHKA *Kcc2/Crh* baseline: N= 8; seizures per day: 1.646 ± 0.566; seizure duration: 41.62 ± 1.856; seizure burden: 2.684 ± 0.789), Gi DREADD treatment substantially inhibited SUDEP incidence in the chronically epileptic *Kcc2/Crh* mice, where only 1 out of 8 mice died of SUDEP compared to 5 out of 13 untreated vIHKA *Kcc2/Crh* mice (Fig. 6C). In fact, the one SUDEP incidence in the chronically epileptic *Kcc2/Crh* group injected with Gi DREADD occurred during the week when CNO was removed from the drinking water. This further suggests that HPA axis dysfunction contributes to SUDEP and that regulating HPA axis activity can reduce the risk of SUDEP.

We also used a pharmacological approach to test whether HPA axis dysfunction in the chronically epileptic *Kcc2/Crh* mice contributed to the increased SUDEP occurrence. Immediately following vIHKA, mice were implanted with a 21-day slow-release pellet of RU486, a synthetic steroid that is a potent glucocorticoid receptor antagonist.^55^ Although RU486 also acts as an antagonist at progesterone receptors^56^ as well as glucocorticoid receptors,^57^ this approach has the benefit of not requiring daily injections which is a major advantage when evaluating the impact of HPA axis on outcome measures. When combined with the Gi DREADD approach described above, we can appropriately interpret these results and draw conclusions from these experiments. There are no significant differences in daily seizure number (Fig. 5C; WT vIHKA: N=6; 1.924 ± 0.542 seizures; *Kcc2/Crh* vIHKA: N= 8; 2.885 ± 0.848 seizures; *Kcc2/Crh* vIHKA+RU486: N= 9; 0.982 ± 0.354) or seizure burden (Fig. 5E; WT vIHKA: N= 6; 4.267 ± 1.15; *Kcc2/Crh* vIHKA: N =8; 6.539 ± 1.984 in vIHKA *Kcc2/Crh*; vIHKA *Kcc2/Crh* +RU486: N=8; 2.293 ± 0.71) in vIHKA *Kcc2/Crh* mice treated with RU486 compared to chronically epileptic WT and *Kcc2/Crh* mice. The vIHKA *Kcc2/Crh* mice treated with RU486 exhibit a significant reduction in the average seizure duration compared to chronically epileptic WT and *Kcc2/Crh* mice (Fig. 5D; WT vIHKA: N= 6; 54.20 ± 3.97 s; *Kcc2/Crh* vIHKA: N= 8; 53.4**9** ± 2.213 s; *Kcc2/Crh* vIHKA+RU486: N= 8; 38.09 ± 2.617 s; one-way ANOVA, \*\**p*< 0.01). Importantly, RU486 treatment prevented SUDEP in the chronically epileptic *Kcc2/Crh* mice, occurring in 0 of the 9 mice tested, compared to 5 out of 13 untreated vIHKA *Kcc2/Crh* (Fig. 6B). These data indicate that HPA axis dysfunction does indeed contribute to SUDEP, and attenuation of seizure induced activation of the HPA axis can reduce SUDEP incidence. Our findings also reveal that the chronically epileptic *Kcc2/Crh* mouse model is a novel, non-genetic model of SUDEP.

### Persistent HPA axis dysfunction in epilepsy may increase SUDEP risk

Data from our preclinical model suggest that HPA axis dysfunction may increase susceptibility to SUDEP, a finding that is incredibly relevant in the clinical setting and could potentially lead to the identification of a biomarker for SUDEP risk. To understand whether our findings translate to the human condition, we ran enzyme linked immunoassays (ELISAs) for corticotropin releasing hormone (CRH) and cortisol in postmortem blood samples collected from PWE with or without suspected SUDEP compared to individuals with no history of epilepsy (samples obtained from the North American SUDEP Registry at NYU Langone Health). Contrary to what we anticipated, we observe a significant decrease in both cortisol and CRH in PWE with suspected SUDEP (cortisol: 1.897 ± 1.271 ng/ml; CRH: 118.1 ± 47.06 ng/ml) compared to either PWE (cortisol: 15.51 ± 5.080 ng/ml; CRH: 270.5 ± 85.71 ng/ml) or individuals with no history of epilepsy (cortisol: 23.04 ± 7.304 ng/ml; CRH: 356.8 ± 154.2 ng/ml) (Fig. 7A and B). This substantial decrease in circulating hormones in the PWE with suspected SUDEP is consistent with HPA axis dysfunction contributing to SUDEP.

**Figure 7:**
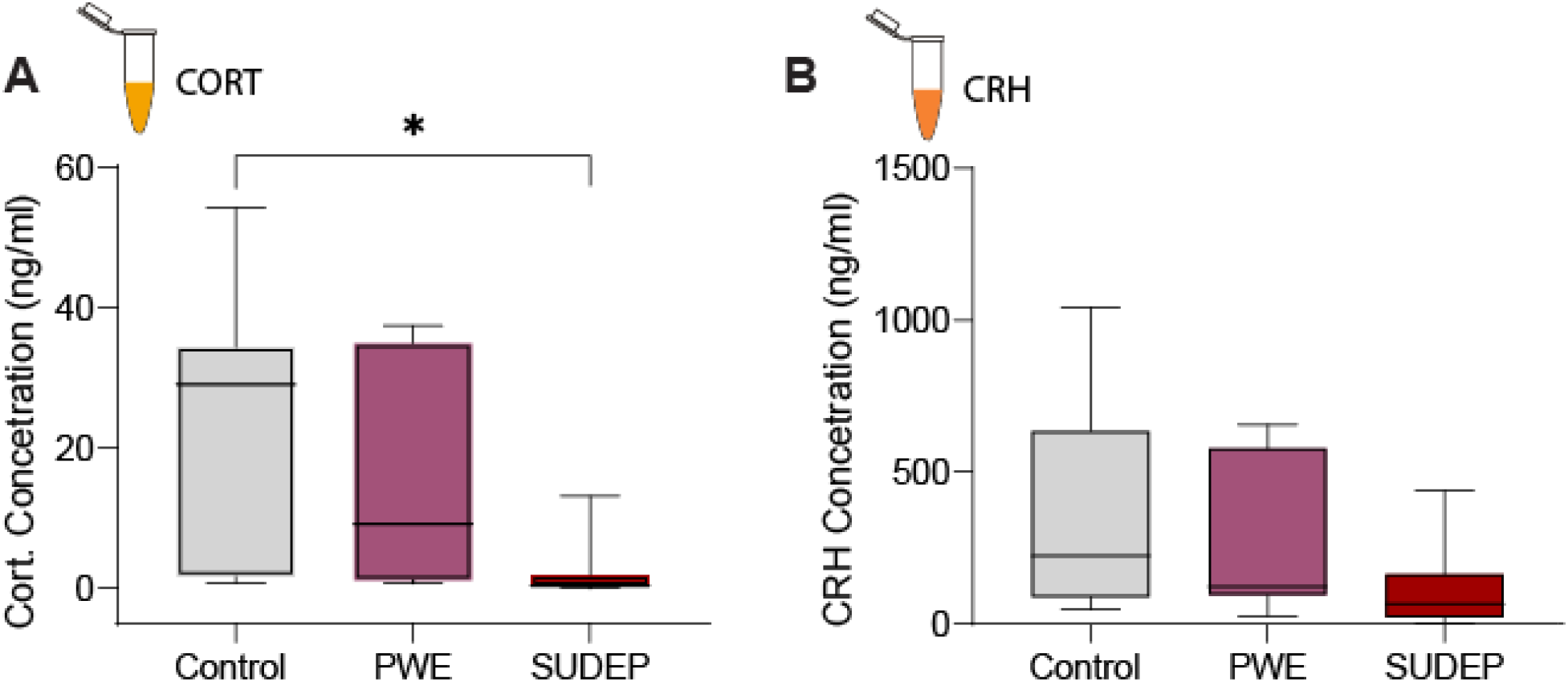
Chronic HPA axis dysfunction may increase SUDEP risk in PWE. **(A-B)** Postmortem analysis of circulating cortisol **(A)** and CRH **(B)** in control subjects, PWE, and subjects who died of SUDEP. Error bars represent SEM. * *p* <0.05. Data was analyzed using a one-way ANOVA. Cort, cortisol; CRH, corticotropin releasing hormone; PWE, person(s) with epilepsy; SUDEP, sudden unexpected death in epilepsy.

We were initially surprised by this finding since we have previously shown that 2 hours after pilocarpine or kainic acid induced SE in mice, there is a significant surge in circulating corticosterone and adrenocorticotropin releasing hormone (ACTH).^2^ However, these data together suggest that following an initial surge of HPA axis activity acutely after a seizure or during the early stage of epileptogenesis, the persistent seizure induced overactivation of the HPA axis leads to its collapse over time, which in turn may substantially increase SUDEP risk. The present data reveals that while the breakdown of the HPA axis occurs in chronic epilepsy, the level of dysfunction over time may be proportional to the intensity of seizure induced activation of the HPA axis, with greater HPA axis dysfunction potentially contributing to increased vulnerability to comorbid mood disorders and greater risk of SUDEP.

## Discussion

Consistent with clinical reports of high incidence rates of anxiety and/or depression in PWE, several studies have shown that chronically epileptic mice exhibit increased anxiety- and depression-like behavior.^17,35,58–60^ We and others have shown that chronically epileptic mice also exhibit an increase in circulating corticosterone levels,^2^ and mitigating HPA axis hyperactivity reduces seizure susceptibility,^24^ neuropathology,^28,29^ and depression-like behavior.^24^ Here, we show that chronically epileptic *Kcc2/Crh* mice have an increased predisposition to anxiety- and depression-like behavior. While there are no significant differences in the aggregated behavioral outcomes in chronically epileptic WT or *Kcc2/Crh* mice, this represents a floor effect in behavioral severity likely due to activation of the HPA axis in both experimental groups. There is, however, a high variability in behavioral outcomes, distributing into resilient and vulnerable populations in which there is an increased vulnerable population in epileptic *Kcc2/Crh* mice compared to WT. Given that seizures activate the HPA axis in WT mice and corticosterone levels are elevated in chronically epileptic WT mice,^2^ further activation of the HPA axis in *Kcc2/Crh* mice may only have a subtle impact on behavioral outcomes, resulting in an increased vulnerability to behavioral deficits associated with epilepsy in *Kcc2/Crh* mice. In support of this interpretation of our data, our previous studies demonstrate that mice with HPA axis hypofunction exhibit significant improvements in behavioral outcomes associated with chronic epilepsy.^24^ Collectively, these studies suggest that HPA axis activation negatively impacts behavioral outcomes associated with epilepsy.

Prior to implementing the vIHKA model^35^ in our *Kcc2/Crh* mice, we tested out the paradigm in a group of C57/Bl6 mice. Like the observations made by Zeidler, et al., we found that chronically epileptic vIHKA C57/Bl6 mice exhibited pronounced differences in OF, LD, FST, SPT, and EPM (data not shown). However, the same pronounced differences are not observed across the behavioral paradigms we tested in the chronically epileptic WT and *Kcc2/Crh* mice. While there are significant differences between the saline injected WT mice and each of the chronically epileptic mouse cohorts in the OF and the FST paradigms, significant differences were not observed in the other paradigms that were tested and there is high variability even within test. Pronounced behavioral differences were not apparent in this study, perhaps due to background strain differences (129/Sv vs. C57Bl6/J). While this is an interesting finding, investigation into the mechanisms contributing to these potential strain differences in behavior is out of the scope of the current study.

Compared to the chronically epileptic WT mice, we observed that a greater proportion of the chronically epileptic *Kcc2/Crh* mice are vulnerable to negative affective states across each of the behavioral paradigms. This led us to hypothesize that there may be separate populations within the *Kcc2/Crh* cohort: a group that underperforms or is more vulnerable to exhibiting negative affective behavior, and another group that may be better at developing coping mechanisms under stressful conditions or are more resilient to anxious- or depressive-like phenotypes. We found that across the behavioral paradigms, the chronically epileptic *Kcc2/Crh* cohort tended to have a greater proportion of mice that were vulnerable to anxiety- or depression-like states compared to the chronically epileptic WT mice. Our data suggests that while chronic epilepsy gives rise to psychiatric comorbidities, increased HPA axis activity in chronic epilepsy may increase predisposition to negative affective states. From a clinical standpoint, screening PWE for changes in circulating cortisol, ACTH, or CRH may provide insight to their risk for developing psychiatric comorbidities, which in turn can dictate a treatment plan to improve their quality of life.

This study does make the unexpected, yet potentially transformative discovery that HPA axis hyperexcitability increases susceptibility to SUDEP. We demonstrate that chronically epileptic mice with exaggerated seizure-induced activation of the HPA axis (*Kcc2/Crh* mouse line ^3^) exhibit increased mortality due to SUDEP, with almost 40% succumbing to SUDEP. To our knowledge, this mouse model represents the first potential environmental link to SUDEP risk. We confirmed that the increased risk of SUDEP in this model is directly related to HPA axis dysfunction since chemogenetic or pharmacological suppression of the HPA axis prevents SUDEP incidence in this model. However, we were also concerned that the SUDEP phenotype may be an artifact of the mouse model with no translational relevance to the human condition. These concerns were dispelled by the evidence of HPA dysfunction in blood samples from PWE that died of suspected SUDEP compared to non-PWE or PWE (without suspected SUDEP) samples. It should be noted that there are other potential variables between the human samples which may contribute to these differences, such as differences in the time of day or time to sample collection, which may indirectly result from the fact that most SUDEP events occur at night.^61–63^. However, cortisol levels are stable over the time of collection,^64,65^ suggesting that this may not be a confounding factor. Thus, these data demonstrate that HPA axis dysfunction may be a novel mechanism contributing to SUDEP and is the first environmental, non-genetic insult to be implicated in the mechanisms contributing to SUDEP.

Stress has been linked to sudden death in people without epilepsy (for review see <u>Lampert</u>, 2014),^66^ primarily due to heart failure.^67^ In fact, it has been estimated that between 20-40% of sudden cardiac deaths are precipitated by stress.^68^ Stress has been shown to increase arrhythmias which has been linked to the increased risk of sudden death.^69^ Although the mechanisms through which stress increases the risk for sudden death is poorly understood, it likely involves the ability of stress to induce changes in autonomic function. ^70,71^

Similar to stress, seizures are also associated with cardiac changes, such as arrhythmias, suggesting that stress should be evaluated as a potential risk factor for SUDEP (for review see Lathers and Schraeder, 2006)^72^. SUDEP is thought to involve autonomic dysfunction^72^ which is tightly regulated by the HPA axis (for review see <u>Ulrich-Lai</u> & <u>Herman</u>, 2009).^73^ Here we demonstrate that HPA axis dysfunction increases SUDEP incidence. These data are the first to link HPA axis dysfunction to SUDEP risk, providing a potential novel mechanism contributing to SUDEP and opening avenues for further mechanistic research into the pathophysiology of SUDEP.

Interestingly, psychiatric illnesses have also been linked to sudden unexpected death unrelated to epilepsy. Psychiatric illnesses are associated with increased morbidity due to numerous factors, including suicide, comorbid alcohol and substance use, and accidents. However, individuals with psychiatric illnesses are also at an increased risk of cardiac sudden death.^74^ For example, depression is associated with an increased risk of cardiovascular disease, coronary heart disease, and cardiac death (for review see <u>Glassman</u>, 2007; <u>Musselman</u>, 1998).^75,76^ The link between psychiatric illnesses and cardiovascular disease has been linked to environment and lifestyle, such as body weight, smoking, and lack of exercise.^77^ However, the exact mechanisms mediating the association between psychiatric illnesses and cardiovascular disease are unresolved (for review see <u>Musselman</u>, 1998).^76^ Relevant to the current study, the HPA axis has also been suggested to mediate the cardiac problems and sudden death,^76,78^ and recently, psychiatric comorbidities associated with epilepsy have been linked to SUDEP^79^

Emerging evidence, including findings presented in this study, demonstrate a link between stress, psychiatric illnesses, sudden death, epilepsy, and SUDEP. Stress activates the HPA axis and is a trigger for psychiatric illnesses. In fact, HPA hyperexcitability is a hallmark feature of depression. Our previous research linked HPA axis dysfunction to comorbid psychiatric illnesses and epilepsy and here we demonstrate a novel mechanistic link to SUDEP. Stress, psychiatric illnesses, and epilepsy have all been linked to cardiovascular disease. Given that the HPA axis influences autonomic and cardiovascular function, future studies will need to examine the mechanistic link between HPA axis dysfunction, autonomic and cardiovascular function, and SUDEP.

A risk factor for SUDEP is high seizure burden and uncontrolled seizures.^80,81^ However, we did not observe significant differences in seizure frequency, duration, or burden between chronically epileptic *Kcc2/Crh* and WT mice. Thus, our evidence does not support an indirect relationship between seizure burden and SUDEP incidence in this model, suggesting a more direct relationship. The lack of effect on seizure frequency may also be due to a ceiling effect since we have shown that suppressing seizure-induced activation of the HPA axis decreases seizure susceptibility and spontaneous seizure frequency in chronically epileptic mice.^24^ Our data suggests that increased HPA axis activity may be a predictor of SUDEP and mitigating its activity may reduce SUDEP susceptibility in PWE. Indeed, we were able to dramatically reduce SUDEP incidence in the chronically epileptic *Kcc2/Crh* mice by attenuating HPA axis function, either through pharmacologically blocking glucocorticoid receptor activity through RU486, or through chemogenetically inhibiting CRH neuron activity using hM4Di-DREADDs. This finding is of incredible clinical relevance and indicates that periodically monitoring circulating stress hormones in PWE may be used as a measure to assess SUDEP risk at any point, and thus may help better guide immediate treatment to prevent SUDEP.

Our use of RU486 in this study was to test the hypothesis that attenuation of the seizure induced hyperactivity of the HPA axis in chronically epileptic *Kcc2/Crh* mice would mitigate SUDEP incidence. Studies have shown that this synthetic steroid potently binds to and antagonizes glucocorticoid receptors.^57^ It is important to note that RU486 can also moderately antagonize progesterone receptors^56^ and can also antagonize androgen receptors^82^ to an even lesser degree and reduced affinity. While it is possible that the RU486 treatment in our study may have acted on all three receptors, studies show that RU486 preferentially binds to glucocorticoid receptors in environments where corticosterone is present in pathologically elevated levels,^83^ which is the case in the chronically epileptic *Kcc2/Crh* mouse model we used in our study. Glucocorticoid receptors are also predominantly activated and maximally saturated over mineralocorticoid receptors in high corticosterone environments,^23,84^ further establishing RU486 specificity to glucocorticoid receptors in our study. While we appreciate the limitations of the pharmacological use of RU486, our evidence suggests that this approach is sufficient to reduce HPA axis activity, and therefore abolish SUDEP incidence in chronically epileptic *Kcc2/Crh* mice. The route of administration of RU486 also has the advantage of not requiring daily injections which are stressful and would confound the results of this study. In combination with our chemogenetic approach to directly evaluate the role of the HPA axis, these studies convincingly demonstrate that HPA axis dysfunction increases SUDEP risk in chronically epileptic *Kcc2/Crh* mice and suppressing HPA hyperexcitability is effective at reducing SUDEP risk.

These findings demonstrate that the *Kcc2/Crh* mouse model is a novel mouse model of SUDEP that the field can use to progress our limited understanding of the mechanisms underlying SUDEP. While some genetic risk factors in PWE increase susceptibility to SUDEP,^33,34^ non-genetic risk factors influencing SUDEP incidence are relatively understudied. Our model suggests that HPA axis dysfunction may be a contributing factor to increased SUDEP risk, suggesting an environmental link to SUDEP risk. A prominent hypothesis in the field is that SUDEP results from cardiac and/or respiratory dysfunction in PWE. Studies have shown that seizures can compromise both cardiac and respiratory ^85–87^ function in PWE. Combined with our previous work showing that seizures alone can activate the HPA axis^2^ and work from others showing that increased HPA axis function independent of seizures contributes to compromised heart function,^88,89^ we hypothesize that increased HPA axis dysfunction in chronic epilepsy can contribute to cardiorespiratory deficits that can predispose PWE to SUDEP.

It is also possible that excessive glucocorticoid signaling in response to repeated and excessive HPA axis activation induces pathological changes, such as cardiac remodeling which may increase the risk to SUDEP.^90,91^ This potential mechanism is supported by evidence of cardiac remodeling in some SUDEP patients.^92–94^ The HPA axis also regulates cardiorespiratory function^91,95,96^ and influences inflammatory processes,^97,98^ both of which could affect SUDEP risk. These potential mechanisms are supported by the fact that SUDEP is a suspected cardiorespiratory event^99,100^ and changes in inflammation have been observed in association with SUDEP.^101,102^ Interestingly, a subpopulation of CRH neurons in the PVN send direct projections to the rostral ventrolateral medulla (RVLM)^103^ and nucleus of the solitary tract (NTS)^104^, which control autonomic and cardiovascular function. Further studies are required to obtain a mechanistic understanding of how HPA axis dysfunction increases vulnerability to SUDEP.

Approximately 60% of SUDEP cases have been reported to occur at night or early morning,^62^ potentially implicating circadian rhythm influence over SUDEP risk. Release or stress hormones, such as cortisol, exhibit a diurnal rhythm,^105,106^ which could influence SUDEP susceptibility. In fact, altered diurnal fluctuations in cortisol have been linked to negative mental and physical health outcomes.^107^ However, to our knowledge, the link between diurnal cortisol levels and SUDEP has not been explored. Here we demonstrate significantly lower levels of circulating stress hormones in postmortem samples from patients with epilepsy that died of confirmed or suspected SUDEP. These findings warrant further exploration of changes in cortisol levels in PWE, particularly those at risk of SUDEP.

Finally, an important caveat to the current study is that we only show data from male mice. The primary question we sought to answer was whether HPA axis hyperactivity in chronic epilepsy contributed to comorbid psychiatric disorders. It is difficult to ignore the fact that globally, females are twice as likely to suffer from depression^108,109^ and present with a more severe symptom profile compared to males, which often renders available treatment options ineffective. With such a substantial global burden, there is a desperate need to understand whether there are sex differences in how HPA axis dysfunction contributes to comorbid depression in chronically epileptic females versus males. Furthermore, sex differences in epilepsy have been well-documented.^81,110,111^ For example, several epidemiological studies document more SUDEP cases among males compared to females.^112,113^ Interestingly, among females suffering from epilepsy, SUDEP incidence rates were reportedly five times higher in those that suffered comorbid psychiatric illness compared to those who did not,^113^ further implicating distinct sex differences in epilepsy and comorbid mood disorders. Ongoing work in our laboratory is focused on understanding how HPA axis dysregulation in chronic epilepsy is regulated between sexes and how potential sex differences in HPA axis dysregulation can influence affective states and SUDEP risk.

## Conclusion

In summary, our work provides new insight into how HPA axis dysfunction may contribute to negative outcomes in epilepsy, including vulnerability to comorbid psychiatric disorders. Importantly, this work demonstrates for the first time that HPA axis dysfunction may play a pathophysiological role in SUDEP. Our data indicates that modulating HPA axis dysfunction in PWE may decrease their risk of developing comorbid psychiatric disorders, mitigate neuropathology, and reduce risk of SUDEP incidence. The current study also highlights a novel mouse model of SUDEP, which will be beneficial to the field for investigating the pathophysiological mechanisms contributing to SUDEP.

## Supplemental Information

The online version contains supplementary material available which will be provided upon request.

## Author Contributions

T.B. and J.M. conceived and designed the study. P.A. and G.L.W. oversaw that data management and analysis pipelines. T.B., P.A., and G.L.W. analyzed the data. T.B., J.M., P.A., and G.L.W. wrote the manuscript. O.D., D.F., and J.L. provided the postmortem human samples used in this study.

## Resource Availability

Requests for information or resources should be directed to the lead contact, Jamie Maguire.

## Materials availability

The data analysis code is available on GitHub. *Kcc2/Crh* mice are available upon request.

## Data and code availability

Raw data used in this study are available from the lead contact upon reasonable request. All custom Python scripts for analysis and visualization are available from the lead contact upon reasonable request. Mobility-mapper for behavioral scoring is available from: https://github.com/researchgrant/mobility-mapper. Seizure detection app is available from: https://github.com/neurosimata/seizy.

## Acknowledgements

T.B., P.A., G.L.W., and J.M. are supported by funding from the National Institutes of Health under award numbers: F31AA028410, R01AA026256, R01NS105628, R01NS102937, R01MH128235, and P50MH122379. O.D. is supported by funding from: NINDS (18-A0-00-1000473, 16-A0-00-006058 / 107311), CDC (26 B 89011), NSF (26 D 70200), and GW Pharma (C19-01030-1012157 / 116548). D.F. is supported by funding from: NINDS (R01 NS109367, R01 NS233102, R01NS123928, 1U44NS121562), NSF (A20 0089 S001), and CDC (6U48DP006396). Human blood samples used in this study were provided by the North American SUDEP Registry (NASR) and funded by Finding A Cure for Epilepsy and Seizures (FACES).

## References

1 Kanner, A. M. Depression in epilepsy: prevalence, clinical semiology, pathogenic mechanisms, and treatment. Biological Psychiatry 54, 388-398-398, doi:10.1016/s0006-3223(03)00469-4 (2003).

2 O’Toole, K. K., Hooper, A., Wakefield, S. & Maguire, J. Seizure-induced disinhibition of the HPA axis increases seizure susceptibility. Epilepsy Res 108, 29–43, doi:10.1016/j.eplepsyres.2013.10.013 (2014).

3 Melon, L. C., Hooper, A., Yang, X., Moss, S. J. & Maguire, J. Inability to suppress the stress-induced activation of the HPA axis during the peripartum period engenders deficits in postpartum behaviors in mice. Psychoneuroendocrinology 90, 182–193, doi:10.1016/j.psyneuen.2017.12.003 (2018).

4 Lambert, M. V. & Robertson, M. M. Depression in epilepsy: etiology, phenomenology, and treatment. Epilepsia 40 Suppl 10, S21–47 (1999).

5 Brandt, C. et al. Prevalence of anxiety disorders in patients with refractory focal epilepsy--a prospective clinic based survey. Epilepsy Behav 17, 259–263, doi:10.1016/j.yebeh.2009.12.009 (2010).

6 Kwon, O.-Y. & Park, S.-P. Depression and Anxiety in People with Epilepsy. Journal of Clinical Neurology 10, 175-188-188, doi:10.3988/jcn.2014.10.3.175 (2014).

7 Mazarati, A. & Sankar, R. Common Mechanisms Underlying Epileptogenesis and the Comorbidities of Epilepsy. Cold Spring Harbor Perspectives in Medicine 6, a022798, doi:10.1101/cshperspect.a022798 (2016).

8 Kanner, A. M. Depression and Epilepsy: A New Perspective on Two Closely Related Disorders. Epilepsy Currents 6, 141-146-146, doi:10.1111/j.1535-7511.2006.00125.x (2006).

9 Christensen, J., Vestergaard, M., Mortensen, P. B., Sidenius, P. & Agerbo, E. Epilepsy and risk of suicide: a population-based case–control study. The Lancet Neurology 6, 693-698-698, doi:10.1016/s1474-4422(07)70175-8 (2007).

10 Kloet, E. R. Hormones and the Stressed Brain. Annals of the New York Academy of Sciences 1018, 1-15-15, doi:10.1196/annals.1296.001 (2004).

11 Swaab, D. F., Bao, A. M. & Lucassen, P. J. The stress system in the human brain in depression and neurodegeneration. Ageing Res Rev 4, 141–194, doi:10.1016/j.arr.2005.03.003 (2005).

12 Gold, P. W., Goodwin, F. K. & Chrousos, G. P. Clinical and Biochemical Manifestations of Depression. The New England Journal of Medicine 319, 413-420-420, doi:10.1056/nejm198808183190706 (1988).

13 Holsboer, F. & Barden, N. Antidepressants and Hypothalamic-Pituitary-Adrenocortical Regulation. Endocrine Reviews 17, 187-205-205, doi:10.1210/edrv-17-2-187 (1996).

14 Arborelius, L., Owens, M. J., Plotsky, P. M. & Nemeroff, C. B. The role of corticotropin-releasing factor in depression and anxiety disorders. Journal of Endocrinology 160, 1-12-12, doi:10.1677/joe.0.1600001 (1999).

15 Sawyer, N. T. & Escayg, A. Stress and epilepsy: multiple models, multiple outcomes. J Clin Neurophysiol 27, 445–452, doi:10.1097/WNP.0b013e3181fe0573 (2010).

16 McKee, H. R. & Privitera, M. D. Stress as a seizure precipitant: Identification, associated factors, and treatment options. Seizure 44, 21-26-26, doi:10.1016/j.seizure.2016.12.009 (2017).

17 Wulsin, A. C. et al. Functional disruption of stress modulatory circuits in a model of temporal lobe epilepsy. PLoS One 13, e0197955, doi:10.1371/journal.pone.0197955 (2018).

18 Culebras, A., Miller, M., Bertram, L. & Koch, J. Differential Response of Growth Hormone, Cortisol, and Prolactin to Seizures and to Stress. Epilepsia 28, 564-570-570, doi:10.1111/j.1528-1157.1987.tb03689.x (1987).

19 Galimberti, C. A. et al. Seizure frequency and cortisol and dehydroepiandrosterone sulfate (DHEAS) levels in women with epilepsy receiving antiepileptic drug treatment. Epilepsia 46, 517–523, doi:10.1111/j.0013-9580.2005.59704.x (2005).

20 Gold, P. W. et al. Responses to corticotropin-releasing hormone in the hypercortisolism of depression and Cushing’s disease. Pathophysiologic and diagnostic implications. N Engl J Med 314, 1329–1335, doi:10.1056/NEJM198605223142101 (1986).

21 Sperling, M. R., Schilling, C. A., Glosser, D., Tracy, J. I. & Asadi-Pooya, A. A. Self-perception of seizure precipitants and their relation to anxiety level, depression, and health locus of control in epilepsy. Seizure 17, 302-307-307, doi:10.1016/j.seizure.2007.09.003 (2008).

22 Neugebauer, R. et al. Stressful Life Events and with Seizure Frequency in Patients Epilepsy. Epilepsia 35, 336-343-343, doi:10.1111/j.1528-1157.1994.tb02441.x (1994).

23 Joёls, M. Stress, the hippocampus, and epilepsy. Epilepsia 50, 586-597-597, doi:10.1111/j.1528-1167.2008.01902.x (2009).

24 Hooper, A., Paracha, R. & Maguire, J. Seizure-induced activation of the HPA axis increases seizure frequency and comorbid depression-like behaviors. Epilepsy Behav 78, 124–133, doi:10.1016/j.yebeh.2017.10.025 (2018).

25 Wulsin, A. C., Solomon, M. B., Privitera, M. D., Danzer, S. C. & Herman, J. P. Hypothalamic-pituitary-adrenocortical axis dysfunction in epilepsy. Physiol Behav 166, 22–31, doi:10.1016/j.physbeh.2016.05.015 (2016).

26 Basu, T., Maguire, J. & Salpekar, J. A. Hypothalamic-pituitary-adrenal axis targets for the treatment of epilepsy. Neurosci Lett 746, 135618, doi:10.1016/j.neulet.2020.135618 (2021).

27 Maguire, J. & Salpekar, J. A. Stress, seizures, and hypothalamic–pituitary–adrenal axis targets for the treatment of epilepsy. Epilepsy & Behavior 26, 352-362-362, doi:10.1016/j.yebeh.2012.09.040 (2013).

28 Wulsin, A. C., Herman, J. P. & Danzer, S. C. RU486 Mitigates Hippocampal Pathology Following Status Epilepticus. Frontiers in Neurology 7, 214, doi:10.3389/fneur.2016.00214 (2016).

29 Wulsin, A. C. et al. The glucocorticoid receptor specific modulator CORT108297 reduces brain pathology following status epilepticus. Exp Neurol 341, 113703, doi:10.1016/j.expneurol.2021.113703 (2021).

30 Herman, J. P., Ostrander, M. M., Mueller, N. K. & Figueiredo, H. Limbic system mechanisms of stress regulation: hypothalamo-pituitary-adrenocortical axis. Progress in neuropsychopharmacology & biological psychiatry 29, 1201–1213, doi:10.1016/j.pnpbp.2005.08.006 (2005).

31 Decavel, C. & Van den Pol, A. N. GABA: a dominant neurotransmitter in the hypothalamus. J Comp Neurol 302, 1019–1037, doi:10.1002/cne.903020423 (1990).

32 Sarkar, J., Wakefield, S., MacKenzie, G., Moss, S. J. & Maguire, J. Neurosteroidogenesis Is Required for the Physiological Response to Stress: Role of Neurosteroid-Sensitive GABAA Receptors. The Journal of Neuroscience 31, 18198-18210-18210, doi:10.1523/JNEUROSCI.2560-11.2011 (2011).

33 Bagnall, R. D., Crompton, D. E. & Semsarian, C. Genetic Basis of Sudden Unexpected Death in Epilepsy. Front Neurol 8, 348, doi:10.3389/fneur.2017.00348 (2017).

34 Coll, M., Oliva, A., Grassi, S., Brugada, R. & Campuzano, O. Update on the Genetic Basis of Sudden Unexpected Death in Epilepsy. Int J Mol Sci 20, doi:10.3390/ijms20081979 (2019).

35 Zeidler, Z. et al. Targeting the Mouse Ventral Hippocampus in the Intrahippocampal Kainic Acid Model of Temporal Lobe Epilepsy. eNeuro 5, doi:10.1523/ENEURO.0158-18.2018 (2018).

36 Melon, L. C., Nasman, J. T., John, A. S., Mbonu, K. & Maguire, J. L. Interneuronal delta-GABAA receptors regulate binge drinking and are necessary for the behavioral effects of early withdrawal. Neuropsychopharmacology 44, 425–434, doi:10.1038/s41386-018-0164-z (2019).

37 Maguire, J. & Mody, I. GABA(A)R plasticity during pregnancy: relevance to postpartum depression. Neuron 59, 207–213, doi:10.1016/j.neuron.2008.06.019 (2008).

38 Mathis, A. et al. DeepLabCut: markerless pose estimation of user-defined body parts with deep learning. Nat Neurosci 21, 1281–1289, doi:10.1038/s41593-018-0209-y (2018).

39 Golde, W. T., Gollobin, P. & Rodriguez, L. L. A rapid, simple, and humane method for submandibular bleeding of mice using a lancet. Lab Anim (NY) 34, 39–43, doi:10.1038/laban1005-39 (2005).

40 Preibisch, S., Saalfeld, S. & Tomancak, P. Globally optimal stitching of tiled 3D microscopic image acquisitions. Bioinformatics 25, 1463–1465, doi:10.1093/bioinformatics/btp184 (2009).

41 Stirling, D. R. et al. CellProfiler 4: improvements in speed, utility and usability. BMC Bioinformatics 22, 433, doi:10.1186/s12859-021-04344-9 (2021).

42 Buckmaster, P. S. & Lew, F. H. Rapamycin suppresses mossy fiber sprouting but not seizure frequency in a mouse model of temporal lobe epilepsy. J Neurosci 31, 2337–2347, doi:10.1523/JNEUROSCI.4852-10.2011 (2011).

43 Parent, J. M., Elliott, R. C., Pleasure, S. J., Barbaro, N. M. & Lowenstein, D. H. Aberrant seizure-induced neurogenesis in experimental temporal lobe epilepsy. Annals of Neurology 59, 81-91-91, doi:10.1002/ana.20699 (2006).

44 Cho, K. O. et al. Aberrant hippocampal neurogenesis contributes to epilepsy and associated cognitive decline. Nat Commun 6, 6606, doi:10.1038/ncomms7606 (2015).

45 Houser, C. R. Granule cell dispersion in the dentate gyrus of humans with temporal lobe epilepsy. Brain Res 535, 195–204, doi:10.1016/0006-8993(90)91601-c (1990).

46 Sutula, T., Cascino, G., Cavazos, J., Parada, I. & Ramirez, L. Mossy fiber synaptic reorganization in the epileptic human temporal lobe. Ann Neurol 26, 321–330, doi:10.1002/ana.410260303 (1989).

47 McEwen, B. S. Stress-induced remodeling of hippocampal CA3 pyramidal neurons. Brain Res 1645, 50–54, doi:10.1016/j.brainres.2015.12.043 (2016).

48 McEwen, B. S. & Magarinos, A. M. Stress Effects on Morphology and Function of the Hippocampus. Annals of the New York Academy of Sciences 821, 271-284-284, doi:10.1111/j.1749-6632.1997.tb48286.x (2017).

49 McEwen, B. S., Nasca, C. & Gray, J. D. Stress Effects on Neuronal Structure: Hippocampus, Amygdala, and Prefrontal Cortex. Neuropsychopharmacology 41, 3, doi:10.1038/npp.2015.171 (2016).

50 de Kloet, E. R., Joels, M. & Holsboer, F. Stress and the brain: from adaptation to disease. Nat Rev Neurosci 6, 463–475, doi:10.1038/nrn1683 (2005).

51 Krugers, H. J., Lucassen, P. J., Karst, H. & Joels, M. Chronic stress effects on hippocampal structure and synaptic function: relevance for depression and normalization by antiglucocorticoid treatment. Front Synaptic Neurosci 2, 24, doi:10.3389/fnsyn.2010.00024 (2010).

52 Porsolt, R. D., Le Pichon, M. & Jalfre, M. Depression: a new animal model sensitive to antidepressant treatments. Nature 266, 730–732, doi:10.1038/266730a0 (1977).

53 Hasegawa, H. & Tomita, H. Assessment of taste disorders in rats by simultaneous study of the two-bottle preference test and abnormal ingestive behavior. Auris Nasus Larynx 13 Suppl 1, S33–41, doi:10.1016/s0385-8146(86)80032-3 (1986).

54 Cathomas, F., Hartmann, M. N., Seifritz, E., Pryce, C. R. & Kaiser, S. The translational study of apathy-an ecological approach. Front Behav Neurosci 9, 241, doi:10.3389/fnbeh.2015.00241 (2015).

55 Peeters, B. W., Ruigt, G. S., Craighead, M. & Kitchener, P. Differential effects of the new glucocorticoid receptor antagonist ORG 34517 and RU486 (mifepristone) on glucocorticoid receptor nuclear translocation in the AtT20 cell line. Ann N Y Acad Sci 1148, 536–541, doi:10.1196/annals.1410.072 (2008).

56 Herrmann, W. et al. [The effects of an antiprogesterone steroid in women: interruption of the menstrual cycle and of early pregnancy]. C R Seances Acad Sci III 294, 933–938 (1982).

57 Peeters, B. W. et al. Glucocorticoid receptor antagonists: new tools to investigate disorders characterized by cortisol hypersecretion. Stress 7, 233–241, doi:10.1080/10253890400019672 (2004).

58 Groticke, I., Hoffmann, K. & Loscher, W. Behavioral alterations in the pilocarpine model of temporal lobe epilepsy in mice. Exp Neurol 207, 329–349, doi:10.1016/j.expneurol.2007.06.021 (2007).

59 Muller, C. J., Groticke, I., Bankstahl, M. & Loscher, W. Behavioral and cognitive alterations, spontaneous seizures, and neuropathology developing after a pilocarpine-induced status epilepticus in C57BL/6 mice. Exp Neurol 219, 284–297, doi:10.1016/j.expneurol.2009.05.035 (2009).

60 Gröticke, I., Hoffmann, K. & Löscher, W. Behavioral alterations in a mouse model of temporal lobe epilepsy induced by intrahippocampal injection of kainate. Experimental Neurology 213, 71-83-83, doi:10.1016/j.expneurol.2008.04.036 (2008).

61 Purnell, B. S., Thijs, R. D. & Buchanan, G. F. Dead in the Night: Sleep-Wake and Time-Of-Day Influences on Sudden Unexpected Death in Epilepsy. Front Neurol 9, 1079, doi:10.3389/fneur.2018.01079 (2018).

62 Lamberts, R. J., Thijs, R. D., Laffan, A., Langan, Y. & Sander, J. W. Sudden unexpected death in epilepsy: people with nocturnal seizures may be at highest risk. Epilepsia 53, 253–257, doi:10.1111/j.1528-1167.2011.03360.x (2012).

63 Devinsky, O., Hesdorffer, D. C., Thurman, D. J., Lhatoo, S. & Richerson, G. Sudden unexpected death in epilepsy: epidemiology, mechanisms, and prevention. Lancet Neurol 15, 1075–1088, doi:10.1016/S1474-4422(16)30158-2 (2016).

64 Garde, A. H. & Hansen, A. M. Long-term stability of salivary cortisol. Scand J Clin Lab Invest 65, 433–436, doi:10.1080/00365510510025773 (2005).

65 Khonmee, J. et al. Effect of time and temperature on stability of progestagens, testosterone and cortisol in Asian elephant blood stored with and without anticoagulant. Conserv Physiol 7, coz031, doi:10.1093/conphys/coz031 (2019).

66 Lampert, R. et al. Triggering of symptomatic atrial fibrillation by negative emotion. J Am Coll Cardiol 64, 1533–1534, doi:10.1016/j.jacc.2014.07.959 (2014).

67 Pignalberi, C., Ricci, R. & Santini, M. [Psychological stress and sudden death]. Ital Heart J Suppl 3, 1011–1021 (2002).

68 Vlastelica, M. Emotional stress as a trigger in sudden cardiac death. Psychiatr Danub 20, 411–414 (2008).

69 Lampert, R. Emotion and sudden cardiac death. Expert Rev Cardiovasc Ther 7, 723–725, doi:10.1586/erc.09.75 (2009).

70 Ziegler, M. G. in Primer on the autonomic nervous system 291–293 (Elsevier, 2012).

71 Ginsberg, J. P. Editorial: Dysregulation of Autonomic Cardiac Control by Traumatic Stress and Anxiety. Front Psychol 7, 945, doi:10.3389/fpsyg.2016.00945 (2016).

72 Lathers, C. M. & Schraeder, P. L. Stress and sudden death. Epilepsy Behav 9, 236–242, doi:10.1016/j.yebeh.2006.06.001 (2006).

73 Ulrich-Lai, Y. M. & Herman, J. P. Neural regulation of endocrine and autonomic stress responses. Nat Rev Neurosci 10, 397–409, doi:10.1038/nrn2647 (2009).

74 Uchida, H. & Suzuki, T. Cardiac Sudden Death in Psychiatric Patients. Can J Psychiatry 60, 203–205, doi:10.1177/070674371506000501 (2015).

75 Glassman, A. H. Depression and cardiovascular comorbidity. Dialogues Clin Neurosci 9, 9–17 (2007).

76 Musselman, D. L., Evans, D. L. & Nemeroff, C. B. The relationship of depression to cardiovascular disease: epidemiology, biology, and treatment. Arch Gen Psychiatry 55, 580–592, doi:10.1001/archpsyc.55.7.580 (1998).

77 Nielsen, R. E., Banner, J. & Jensen, S. E. Cardiovascular disease in patients with severe mental illness. Nat Rev Cardiol 18, 136–145, doi:10.1038/s41569-020-00463-7 (2021).

78 Jokinen, J. & Nordstrom, P. HPA axis hyperactivity and cardiovascular mortality in mood disorder inpatients. J Affect Disord 116, 88–92, doi:10.1016/j.jad.2008.10.025 (2009).

79 Tao, G. et al. Association Between Psychiatric Comorbidities and Mortality in Epilepsy. Neurol Clin Pract 11, 429–437, doi:10.1212/CPJ.0000000000001114 (2021).

80 Tomson, T., Walczak, T., Sillanpaa, M. & Sander, J. W. Sudden unexpected death in epilepsy: a review of incidence and risk factors. Epilepsia 46 Suppl 11, 54–61, doi:10.1111/j.1528-1167.2005.00411.x (2005).

81 Hesdorffer, D. C. et al. Combined analysis of risk factors for SUDEP. Epilepsia 52, 1150–1159, doi:10.1111/j.1528-1167.2010.02952.x (2011).

82 Song, L. N., Coghlan, M. & Gelmann, E. P. Antiandrogen effects of mifepristone on coactivator and corepressor interactions with the androgen receptor. Mol Endocrinol 18, 70–85, doi:10.1210/me.2003-0189 (2004).

83 Spitz, I. M. & Bardin, C. W. Mifepristone (RU 486)--a modulator of progestin and glucocorticoid action. N Engl J Med 329, 404–412, doi:10.1056/NEJM199308053290607 (1993).

84 Tasker, J. G., Di, S. & Malcher-Lopes, R. Rapid Glucocorticoid Signaling via Membrane-Associated Receptors. Endocrinology 147, 5549-5556-5556, doi:10.1210/en.2006-0981 (2006).

85 Devinsky, O. Effects of Seizures on Autonomic and Cardiovascular Function. Epilepsy Curr 4, 43–46, doi:10.1111/j.1535-7597.2004.42001.x (2004).

86 Dlouhy, B. J. et al. Breathing Inhibited When Seizures Spread to the Amygdala and upon Amygdala Stimulation. J Neurosci 35, 10281–10289, doi:10.1523/JNEUROSCI.0888-15.2015 (2015).

87 Bateman, L. M., Spitz, M. & Seyal, M. Ictal hypoventilation contributes to cardiac arrhythmia and SUDEP: report on two deaths in video-EEG-monitored patients. Epilepsia 51, 916–920, doi:10.1111/j.1528-1167.2009.02513.x (2010).

88 Kadmiel, M. & Cidlowski, J. A. Glucocorticoid receptor signaling in health and disease. Trends Pharmacol Sci 34, 518–530, doi:10.1016/j.tips.2013.07.003 (2013).

89 Pimenta, E., Wolley, M. & Stowasser, M. Adverse cardiovascular outcomes of corticosteroid excess. Endocrinology 153, 5137–5142, doi:10.1210/en.2012-1573 (2012).

90 Popovic, D. et al. How does stress possibly affect cardiac remodeling? Peptides 57, 20–30, doi:10.1016/j.peptides.2014.04.006 (2014).

91 Burford, N. G., Webster, N. A. & Cruz-Topete, D. Hypothalamic-Pituitary-Adrenal Axis Modulation of Glucocorticoids in the Cardiovascular System. Int J Mol Sci 18, doi:10.3390/ijms18102150 (2017).

92 Surges, R., Thijs, R. D., Tan, H. L. & Sander, J. W. Sudden unexpected death in epilepsy: risk factors and potential pathomechanisms. Nat Rev Neurol 5, 492–504, doi:10.1038/nrneurol.2009.118 (2009).

93 Devinsky, O. Sudden, unexpected death in epilepsy. N Engl J Med 365, 1801–1811, doi:10.1056/NEJMra1010481 (2011).

94 Li, M. C. H., O’Brien, T. J., Todaro, M. & Powell, K. L. Acquired cardiac channelopathies in epilepsy: Evidence, mechanisms, and clinical significance. Epilepsia 60, 1753–1767, doi:10.1111/epi.16301 (2019).

95 Carney, R. M., Freedland, K. E. & Sheps, D. S. Depression is a risk factor for mortality in coronary heart disease. Psychosom Med 66, 799–801, doi:10.1097/01.psy.0000146795.38162.b1 (2004).

96 Carney, R. M. et al. Depression, heart rate variability, and acute myocardial infarction. Circulation 104, 2024–2028, doi:10.1161/hc4201.097834 (2001).

97 O’Connor, T. M., O’Halloran, D. J. & Shanahan, F. The stress response and the hypothalamic-pituitary-adrenal axis: from molecule to melancholia. QJM 93, 323–333, doi:10.1093/qjmed/93.6.323 (2000).

98 Silverman, M. N. & Sternberg, E. M. Glucocorticoid regulation of inflammation and its functional correlates: from HPA axis to glucocorticoid receptor dysfunction. Ann N Y Acad Sci 1261, 55–63, doi:10.1111/j.1749-6632.2012.06633.x (2012).

99 Lee, J. & Devinsky, O. The role of autonomic dysfunction in sudden unexplained death in epilepsy patients. Rev Neurol Dis 2, 61–69 (2005).

100 Allen, L. A. et al. Dysfunctional Brain Networking among Autonomic Regulatory Structures in Temporal Lobe Epilepsy Patients at High Risk of Sudden Unexpected Death in Epilepsy. Front Neurol 8, 544, doi:10.3389/fneur.2017.00544 (2017).

101 Michalak, Z., Obari, D., Ellis, M., Thom, M. & Sisodiya, S. M. Neuropathology of SUDEP: Role of inflammation, blood-brain barrier impairment, and hypoxia. Neurology 88, 551–561, doi:10.1212/WNL.0000000000003584 (2017).

102 Somani, A., El-Hachami, H., Patodia, S., Sisodiya, S. & Thom, M. Regional microglial populations in central autonomic brain regions in SUDEP. Epilepsia 62, 1318–1328, doi:10.1111/epi.16904 (2021).

103 Dempsey, B. et al. Mapping and Analysis of the Connectome of Sympathetic Premotor Neurons in the Rostral Ventrolateral Medulla of the Rat Using a Volumetric Brain Atlas. Front Neural Circuits 11, 9, doi:10.3389/fncir.2017.00009 (2017).

104 Wang, L. A., Nguyen, D. H. & Mifflin, S. W. Corticotropin-releasing hormone projections from the paraventricular nucleus of the hypothalamus to the nucleus of the solitary tract increase blood pressure. J Neurophysiol 121, 602–608, doi:10.1152/jn.00623.2018 (2019).

105 Veldhuis, J. D., Iranmanesh, A., Lizarralde, G. & Johnson, M. L. Amplitude modulation of a burstlike mode of cortisol secretion subserves the circadian glucocorticoid rhythm. Am J Physiol 257, E6–14, doi:10.1152/ajpendo.1989.257.1.E6 (1989).

106 Veldhuis, J. D., Iranmanesh, A., Johnson, M. L. & Lizarralde, G. Amplitude, but not frequency, modulation of adrenocorticotropin secretory bursts gives rise to the nyctohemeral rhythm of the corticotropic axis in man. J Clin Endocrinol Metab 71, 452–463, doi:10.1210/jcem-71-2-452 (1990).

107 Adam, E. K. et al. Diurnal cortisol slopes and mental and physical health outcomes: A systematic review and meta-analysis. Psychoneuroendocrinology 83, 25–41, doi:10.1016/j.psyneuen.2017.05.018 (2017).

108 Whiteford, H. A. et al. Global burden of disease attributable to mental and substance use disorders: findings from the Global Burden of Disease Study 2010. Lancet 382, 1575–1586, doi:10.1016/S0140-6736(13)61611-6 (2013).

109 Kessler, R. C., McGonagle, K. A., Swartz, M., Blazer, D. G. & Nelson, C. B. Sex and depression in the National Comorbidity Survey. I: Lifetime prevalence, chronicity and recurrence. J Affect Disord 29, 85–96, doi:10.1016/0165-0327(93)90026-g (1993).

110 Scharfman, H. E. & MacLusky, N. J. Sex differences in the neurobiology of epilepsy: a preclinical perspective. Neurobiology of disease 72 Pt B, 180–192, doi:10.1016/j.nbd.2014.07.004 (2014).

111 McHugh, J. C. & Delanty, N. Epidemiology and classification of epilepsy: gender comparisons. Int Rev Neurobiol 83, 11–26, doi:10.1016/S0074-7742(08)00002-0 (2008).

112 Verducci, C. et al. SUDEP in the North American SUDEP Registry: The full spectrum of epilepsies. Neurology 93, e227–e236, doi:10.1212/WNL.0000000000007778 (2019).

113 Sveinsson, O., Andersson, T., Carlsson, S. & Tomson, T. The incidence of SUDEP: A nationwide population-based cohort study. Neurology 89, 170–177, doi:10.1212/WNL.0000000000004094 (2017).

